# Propagating population activity patterns during spontaneous slow waves in the thalamus of rodents

**DOI:** 10.1101/2023.08.31.555472

**Authors:** Csaba Horváth, István Ulbert, Richárd Fiáth

**Affiliations:** Institute of Cognitive Neuroscience and Psychology, Research Centre for Natural Sciences, Eötvös Loránd Research Network, Budapest, Hungary; János Szentágothai Doctoral School of Neurosciences, Semmelweis University, Budapest, Hungary; Faculty of Information Technology and Bionics, Pázmány Péter Catholic University, Budapest, Hungary

**Keywords:** slow waves, slow oscillation, up-state, thalamus, propagating activity, multiunit activity

## Abstract

Slow waves (SWs) represent the most prominent electrophysiological events in the thalamocortical system under anesthesia and during deep sleep. Recent studies have revealed that SWs have complex spatiotemporal dynamics and propagate across neocortical regions. However, it is still unclear whether neuronal activity in the thalamus exhibits similar propagation properties during SWs. Here, we report propagating population activity in the thalamus of ketamine/xylazine-anesthetized rats and mice visualized by high-density silicon probe recordings. In both rodent species, propagation of spontaneous thalamic activity during up-states was most frequently observed in dorsal thalamic nuclei such as the higher order posterior (Po), lateral posterior (LP) or laterodorsal (LD) nuclei. The preferred direction of thalamic activity spreading was along the dorsoventral axis, with over half of the up-states exhibiting a gradual propagation in the ventral-to-dorsal direction. Furthermore, simultaneous neocortical and thalamic recordings demonstrated that there is a weak but noticeable interrelation between propagation patterns observed during cortical up-states and those displayed by thalamic population activity. In addition, using chronically implanted silicon probes, we detected propagating activity patterns in the thalamus of naturally sleeping rats during slow-wave sleep. However, in comparison to propagating up-states observed under anesthesia, these propagating patterns were characterized by a reduced rate of occurrence and a faster propagation speed. Our findings suggest that the propagation of spontaneous population activity is an intrinsic property of the thalamocortical network during synchronized brain states such as deep sleep or anesthesia. Additionally, our data implies that the neocortex may have partial control over the formation of propagation patterns within the dorsal thalamus.

## 1. Introduction

Slow waves (SWs) are the predominant brain rhythm observed under anesthesia and during slow-wave sleep (i.e., the deepest stage of non-rapid eye movement sleep; Achermann and Borbely, 1997; Steriade et al., 1993a; Steriade et al., 1993c). These high amplitude, low frequency (∼1 Hz) fluctuations reflect the concerted and near-synchronous activity of large populations of neurons within the thalamocortical network. At the cellular level, the membrane potential of these neurons rhythmically alternates between a hyperpolarized and a depolarized state (Steriade et al., 1993a; Steriade et al., 1993c; Wilson and Kawaguchi, 1996). The latter phase, known as the up-state (or ON period), is characterized by intense synaptic activity and elevated neuronal firing that lasts for several hundred milliseconds, whereas during hyperpolarized periods (referred to as down-states or OFF periods) virtually all neurons cease to fire for a similar period of time.

Over the past two decades, numerous properties of this rhythmic slow oscillation have been described in studies conducted in various animal models using a wide range of in vitro (Capone et al., 2019; Sanchez-Vives and McCormick, 2000; Wester and Contreras, 2012) and in vivo techniques (Chauvette et al., 2011; Chauvette et al., 2010; Dasilva et al., 2021; Hangya et al., 2011; Mohajerani et al., 2010; Sakata and Harris, 2009; Stroh et al., 2013; Vyazovskiy et al., 2009). These findings have provided a deeper insight into the cellular and network mechanisms underlying the generation and complex spatiotemporal dynamics of this phenomenon. For instance, animal studies have provided compelling evidence that the neocortex is the main generator of SWs (Steriade et al., 1993a; Steriade et al., 1993b, c; Timofeev et al., 2000; Timofeev and Steriade, 1996), with cortical layer 5 playing a prominent role in the ignition and horizontal spreading of up-states, at least in animal models (Beltramo et al., 2013; Chauvette et al., 2010; Fiath et al., 2016b; Sakata and Harris, 2009; Sanchez-Vives and McCormick, 2000; Stroh et al., 2013; Wester and Contreras, 2012). However, recent findings suggest that the thalamus also has an essential role in shaping and maintaining neuronal activity during slow waves (Crunelli et al., 2015; Crunelli and Hughes, 2010; David et al., 2013; Lemieux et al., 2014). In addition to animal studies, human research investigating the characteristics of SWs in intracortical and electroencephalography (EEG) recordings has also contributed invaluable knowledge to the field (Achermann and Borbely, 1997; Csercsa et al., 2010; Hangya et al., 2011; Massimini et al., 2004; Murphy et al., 2009; Riedner et al., 2007).

One of the hypothesized fundamental functions of SWs is the consolidation of hippocampus-dependent episodic memories. This hypothesis is supported by recent findings demonstrating the ability of this rhythm to orchestrate the communication between the hippocampus and the thalamocortical network by temporally grouping various faster brain oscillations (e.g., thalamic spindles) and hippocampal sharp-wave ripples (Born et al., 2006; Eschenko et al., 2006; Marshall et al., 2006; Molle and Born, 2011; Ngo et al., 2013; Rasch and Born, 2013; Sirota et al., 2003; Tononi and Cirelli, 2014). Furthermore, accumulating evidence suggests that SWs are also implicated in the restorative function of sleep. For example, high slow-wave activity may effectively drive the clearance of waste products from the brain tissue via the glymphatic system (Hablitz et al., 2019; Xie et al., 2013), whereas down-states characterized by neuronal inactivity might provide a temporal window for cellular rest and maintenance (Vyazovskiy and Harris, 2013).

The development and application of advanced electrophysiological recording methods that provide high spatial resolution concurrently with large cortical coverage has led to the recent discovery of the propagating nature and complex spatiotemporal dynamics of SWs. The traveling of cortical SWs has been observed in humans as well as in various animal species using high-density EEG (Massimini et al., 2004; Murphy et al., 2009; Riedner et al., 2007), electrocorticography (ECoG; Dasilva et al., 2021; Hangya et al., 2011; Pazienti et al., 2022) and multielectrode array recordings (Fucke et al., 2011; Greenberg and Dickson, 2013; Luczak et al., 2007; Sakata and Harris, 2009; Sanchez-Vives and McCormick, 2000). Moreover, cortical SW propagation has also been detected in optical signals obtained with mesoscopic optical imaging methods, such as calcium imaging or voltage-sensitive dye imaging (Afrashteh et al., 2021; Brier et al., 2019; Ferezou et al., 2006; Greenberg et al., 2018; Liang et al., 2021; MacDowell and Buschman, 2020; Mohajerani et al., 2013; Mohajerani et al., 2010; Niethard et al., 2018; Petersen et al., 2003; Reyes-Puerta et al., 2016; Stroh et al., 2013; Takagaki et al., 2008; Vanni et al., 2017; Xiao et al., 2017; Xu et al., 2007).

In humans, for instance, individual slow waves initiate most frequently in an anterior cortical region, although they can start at any point of the cortex, and then typically travel in the posterior direction (Massimini et al., 2004). On a finer spatial scale, however, SWs can be characterized by more complex propagation patterns (Hangya et al., 2011). In rodents, during anesthesia or quiet wakefulness, spontaneous cortical slow waves predominantly propagate along the anterior-posterior axis, either, similarly to what was observed in humans, moving from anterior cortical areas towards posterior regions (anterior-to-posterior or front-to-back propagation), or from posterior parts of the cortex to anterior areas (posterior-to-anterior or back-to-front propagation; Brier et al., 2019; Dasilva et al., 2021; Greenberg et al., 2018; Greenberg and Dickson, 2013; Liang et al., 2021; Matsui et al., 2016; Niethard et al., 2018; Ruiz-Mejias et al., 2011; Sheroziya and Timofeev, 2014; Vanni et al., 2017). Furthermore, several studies have described unique and recurring spatiotemporal patterns of cortical activity, referred to as ‘motifs’, where neuronal activity starts in specific cortical areas and then travels along well-defined trajectories (Afrashteh et al., 2021; MacDowell and Buschman, 2020; Mohajerani et al., 2013).

The majority of related research has focused on investigating the propagation patterns of SWs within the neocortex, while the spatiotemporal activity patterns generated in the thalamus, the other brain structure contributing to the generation of slow waves, have received little attention so far (Sheroziya and Timofeev, 2014; Slezia et al., 2011; Ujma et al., 2022; Ushimaru et al., 2012). One plausible explanation for this limited interest is that the thalamus lies deep within the brain, several millimeters below the neocortex, making it challenging to access, even with invasive methods. Optical imaging techniques are generally unsuitable for capturing thalamic activity, while most penetrating probes used for electrophysiological measurements lack the necessary spatial resolution and spatial coverage to provide a detailed view of the complex spatiotemporal dynamics of thalamic SWs across multiple, small thalamic nuclei. However, this situation may change with recently developed high-electrode-density silicon-based probes (such as the Neuropixels probe; Jun et al., 2017; Steinmetz et al., 2021), which offer a great opportunity to investigate the electrical activity of the thalamus at a finer spatial scale.

In this study, our aim was to investigate spontaneously occurring thalamic up-states in ketamine/xylazine-anesthetized rats and mice, with the goal of revealing propagating population activity within the thalamus or within specific thalamic nuclei. To accomplish this, we used single- and multi-shank high-density silicon probes to collect neuronal activity simultaneously in multiple thalamic nuclei, then we assessed the properties of propagation patterns detected during up-states. Furthermore, with simultaneous cortical and thalamic silicon probe recordings in rats, we also examined the relationship between propagation patterns emerging in these two interconnected brain areas. Moreover, in chronically implanted, naturally sleeping rats, we investigated spiking activity in thalamic nuclei where propagating slow waves had been most frequently observed under anesthesia. We then compared the properties of propagating events identified during non-rapid eye movement sleep with the properties of propagating thalamic up-states observed under anesthesia.

## 2. Materials and Methods

### 2.1. Animals

All experiments were performed in accordance with the EC Council Directive of September 22, 2010 (2010/63/EU), and all procedures were reviewed and approved by the Animal Care Committee of the Research Centre for Natural Sciences and by the National Food Chain Safety Office of Hungary (license number: PE/EA/1261-7/2020). In acute experiments, thalamocortical recordings were obtained from adult Wistar rats (n = 34; weight: 285.22 ± 81.21 g, mean ± SD; n = 19 female) and adult C57BL/6J mice (n = 9; weight: 26.33 ± 7.00 g; n = 4 female). Two adult Wistar rats (259 g, female; 476 g, male) were used for chronic recordings. All experiments were performed during the subjective nighttime state of the animals.

### 2.2. Acute in vivo experiments in anesthetized animals

#### 2.2.1. Animal surgery

For both rats and mice, anesthesia was induced by administering a mixture of ketamine (75 mg/kg body weight for rats and 100 mg/kg for mice) and xylazine (10 mg/kg body weight for both rats and mice) intraperitoneally or intramuscularly. To maintain the deep level of anesthesia during experiments and to sustain a stable slow-wave activity, additional doses of ketamine/xylazine were regularly injected during surgery and subsequent electrophysiological recordings (1 – 2 injections/hour). The physiological body temperature of the animals was maintained by a homeothermic heating pad placed under them. The temperature of the heating pad containing internal temperature sensors was controlled by a temperature controller unit (Supertech, Pécs, Hungary). After the animals reached the level of surgical anesthesia, they were placed in a stereotaxic frame (David Kopf Instruments, Tujunga, CA, USA), and then the skin and connective tissue were removed to expose their skull. Next, using a dental drill, a circular craniotomy with a diameter of about 3 mm was prepared over the targeted thalamic regions of the left hemisphere. In rats, the coordinates of the examined thalamic nuclei ranged from approximately -1.5 mm to -6 mm along the anterior-posterior (AP) axis (from -1 mm to -3 mm in mice), and from approximately 0.4 mm to 4 mm along the medial-lateral (ML) axis (from 0.6 mm to 2.7 mm in mice), with respect to the bregma (Paxinos and Franklin, 2004; Paxinos and Watson, 2007). To minimize brain dimpling during the insertion of silicon probes, small incisions were carefully made in the dura mater over the target sites with a 34-gauge needle. Single- or multi-shank silicon probes were mounted on motorized stereotaxic micromanipulators (Robot Stereotaxic, Neurostar, Tübingen, Germany) and driven into the brain tissue at a slow insertion speed (∼2 μm/s) to a depth of about 5 – 8 mm in rats and 3 – 4.5 mm in mice to reach the thalamus. Slow insertion of silicon probes reduces tissue damage which may result in neuronal recordings with an improved quality (Fiath et al., 2019). During thalamic activity measurements, room temperature physiological saline solution was regularly dripped into the craniotomy to prevent dehydration of the brain tissue. To increase the number of thalamic locations examined in a single animal, consecutive insertions (2 – 4) were performed with the same or another (unused) silicon probe of the same type. A distance of at least 500 µm was kept between the insertions to avoid recording from the proximity of thalamic tissue damaged by one of the previous penetrations. A stainless steel needle inserted into the nuchal muscle of animals served as an external reference electrode and for grounding. For post-mortem histological verification of the recording locations, the silicon shank(s) of the probe were coated with red-fluorescent dye 1,1-dioctadecyl-3,3,3,3-tetramethylindocarbocyanine perchlorate (DiI, D-282, ∼10% in ethanol, Thermo Fisher Scientific, Waltham, MA, USA) prior to insertion. At the end of the experiment, the silicon probes were withdrawn and cleaned in 1% Tergazyme solution (Sigma-Aldrich, St. Louis, MO, USA) for at least 30 minutes, then carefully rinsed with distilled water.

#### 2.2.2. Electrophysiological recordings from the thalamus

Two types of silicon-based probes were used to record spontaneous thalamic activity in anesthetized rodents. To determine whether activity propagation in the thalamus occurs on a larger spatial scale, we used multi-shank probes that provide a larger thalamic coverage, and allow mapping the simultaneous activity of multiple nuclei (A8×8-10mm-200-200-177 probes with 64 recording sites for rats, and A8×16-Edge-5mm-100-200-177 probes with 128 recording sites for mice; NeuroNexus Technologies, Ann Arbor, MI, USA). These probes contain eight silicon shanks with an inter-shank spacing of 200 μm, and multiple, equidistantly spaced recording sites (electrodes) on each shank (8 sites/shank with 200 μm inter-electrode distance for rats; 16 sites/shank with 100 μm inter-electrode distance for mice). Because the dorsoventral coverage of these probes is only about 1.5 mm and the dorsoventral expanse of the thalamus is about 3 – 3.5 mm in rats and 1.5 – 2 mm in mice, we recorded sequentially from multiple (2-3), overlapping thalamic depths, starting from the most dorsal position and advancing the probe ventrally in 500 – 700 μm steps after each recording. Wideband thalamic activity (0.1 – 7500 Hz) was recorded using an Intan RHD-2000 electrophysiological recording system (Intan Technologies, Los Angeles, CA, USA) with a sampling frequency of 20 kHz/channel and a resolution of 16 bits. A single 64-channel Intan amplifier board was used to record from rats and a 128-channel amplifier board was used to record from mice. The recording system was connected to a laptop via USB 2.0. The collected continuous thalamic recordings were saved to a network attached storage device for later data analysis. From 14 rats, 82 spontaneous recordings were acquired with multi-shank silicon probes with an average duration of about 30 minutes, while 18 spontaneous recordings were obtained from 5 mice (see Tables 1 and 2 for more details). In 10 out of 14 rats (and in all mice), the orientation of the probe shanks was parallel to the medial-lateral axis, while in the remaining four animals the probe shanks were aligned along the anterior-posterior axis.

**Table 1.** Summary statistics of recordings obtained in anesthetized rats.

| Rat | Multi-shank recordings | Neuropixels recordings |
| --- | --- | --- |
| No. of animals | 14 | 20 |
| No. of penetrations | 40 | 46 |
| Average number of penetrations per animal | $2.86 \pm 0.86$ | $2.30 \pm 1.13$ |
| No. of recordings | 82 | 46 |
| Average number of recordings per animal | $5.86 \pm 2.03$ | $2.30 \pm 1.13$ |
| Average duration of recordings (min) | $29.93 \pm 6.50$ | $56.44 \pm 17.54$ |

**Table 2.**
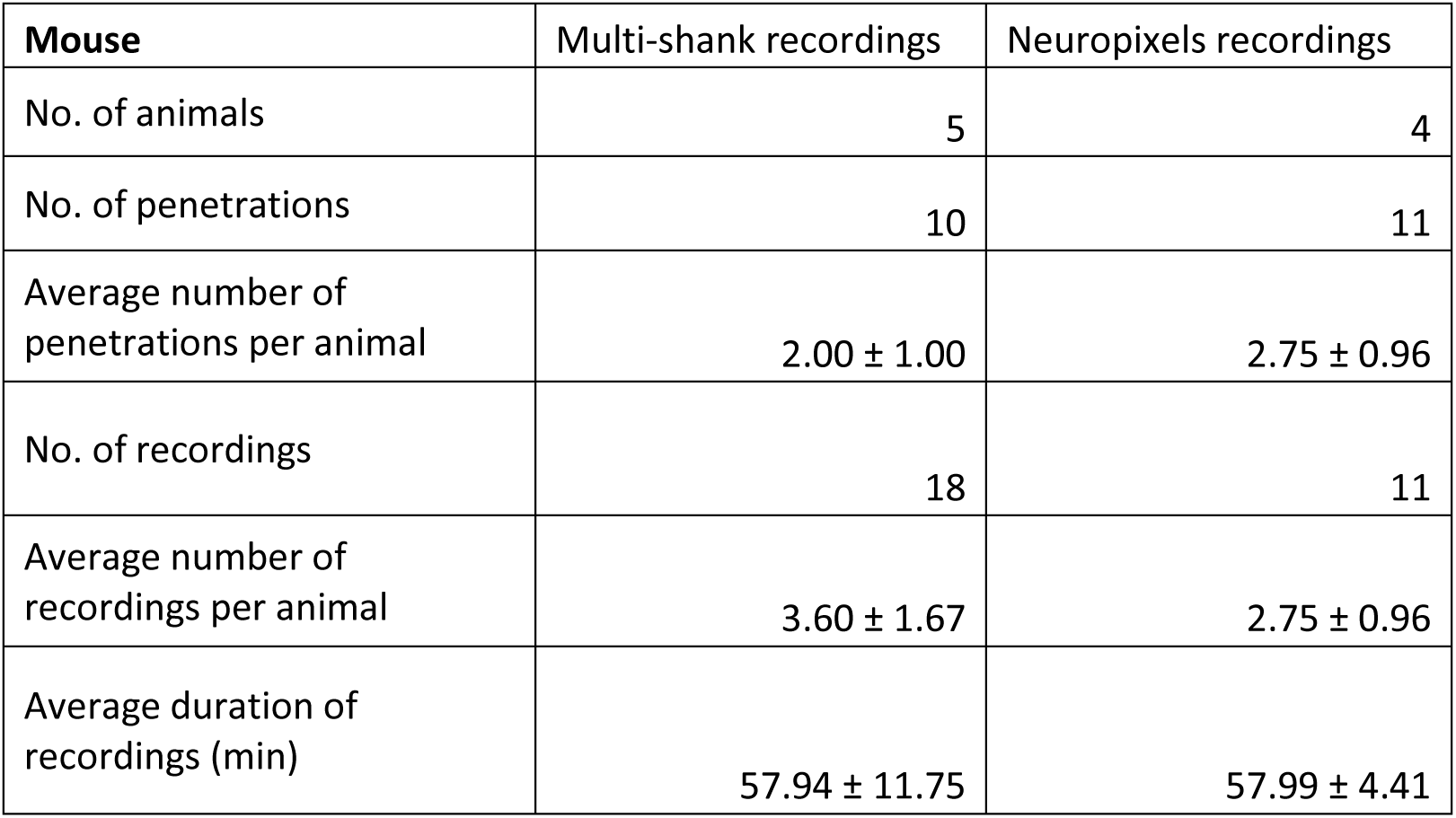
Summary statistics of recordings obtained in anesthetized mice.

To record thalamic activity on a finer spatial scale, we used high-density single-shank Neuropixels 1.0 probes (IMEC, Leuven, Belgium) with 960 recording sites (electrodes) from which 384 sites can be selected for simultaneous recording. The Neuropixels recording system and a PXIe-1071 chassis with an MXI-Express interface (National Instruments, Austin, TX, USA) were used for data acquisition. Spiking activity from the thalamus was recorded at a sampling rate of 30 kHz/channel and with a resolution of 10 bits (amplifier gain: 500; high-pass filter cutoff frequency: 300 Hz), using the SpikeGLX software package (https://billkarsh.github.io/SpikeGLX/). Local field potentials were acquired in separate files at a sampling rate of 2.5 kHz/channel (amplifier gain: 250). From a single animal we typically collected about two to four hours of neuronal data from two to three thalamic insertions. Recording sites located closest to the probe tip (bank 0) were used for data acquisition. From 20 rats, 46 spontaneous recordings were acquired with Neuropixels high-density probes (5 recordings were excluded from further analysis due to low signal quality or irregular slow-wave activity), while 11 recordings were obtained from 4 mice (see Tables 1 and 2 for more details). The duration of each recording was approximately one hour.

#### 2.2.3. Concurrent electrophysiological recordings from the thalamus and the neocortex

In a subset of rat experiments (n = 5 animals), besides the thalamically positioned Neuropixels probe, a multi-shank silicon probe (NeuroNexus Buzsaki64) was inserted into the neocortex to simultaneously record spontaneous population activity from cortical and thalamic regions. This specific multi-shank probe consists of eight shanks with an inter-shank distance of 200 μm and eight closely packed sites located at the tip of each shank, arranged in a V-shape. The multi-shank silicon probe was inserted to a depth of 1.1 mm from the surface of the cortex to record the neuronal activity of layer 5. The shanks of the probe were aligned along the anterior-posterior axis (providing a cortical coverage of ∼1.5 mm along the AP axis), with the first shank located in the most anterior position. The probe was implanted at a lateral inclination of 15° from the vertical (relative to the sagittal plane), so that the shanks were perpendicular to cortical layers. Most of the probe shanks were located in the somatosensory cortex but some of the posterior shanks may have recorded the activity of the parietal association cortex (approximate position of the first shank, AP: -1.5 mm; ML: +4 mm from the bregma). The Neuropixels probe was inserted to a dorsoventral depth of 7 - 7.5 mm (target coordinates, AP: -3 mm; ML: +2 mm), into the thalamic region incorporating first order somatosensory (e.g., the ventral posteromedial nucleus, VPM) as well as higher order nuclei (e.g., the posterior nucleus, Po; or the laterodorsal nucleus, LD). Brain electrical activity was acquired simultaneously with the two recording systems (Intan-based and Neuropixels systems), at a sampling rate of 30 kHz (other recording parameters were the same as described above). The two recording systems were synchronized by starting cortical and thalamic recordings simultaneously with TTL (transistor-transistor logic) pulses generated by a PulsePal device (Sanders and Kepecs, 2014).

### 2.3. Chronic in vivo experiments in freely moving, naturally sleeping rats

#### 2.3.1. Animal surgery

Most of the equipment (e.g., stereotaxic frame, heating pad) used for chronic probe implantations was the same as that used for the acute experiments. For the surgical procedure, we essentially followed the steps described in van Daal et al. (2021). Prior to probe implantation, a reusable 3D-printed single-probe fixture (ATLAS Neuroengineering, Leuven, Belgium) consisting of three main parts was prepared (van Daal et al., 2021). The main body of the fixture contains a single-shank Neuropixels 1.0 silicon probe and a headstage previously fixed to the device. The skull connector is designed to secure the fixture to the skull, while the protective cover prevents damage to the electronic components. Following the assembly of the 3D-printed fixture, the silicon shank of the Neuropixels probe was coated with fluorescent DiI to confirm the recording location in the thalamus post-mortem.

Surgical tools were sterilized before the implantation procedure. Adult Wistar rats (n = 2) were anesthetized with ketamine/xylazine in the same way as described above for the acute surgery. After the animal reached the surgical level of anesthesia, 0.1 ml meloxicam (Dopharma Research, Raamsdonksveer, Netherlands) was administered subcutaneously (1 mg/kg), then the rat was placed in the stereotaxic frame. The physiological body temperature of the animal was maintained by the homeothermic heating pad. Ophthalmic gel (Corneregel, Dr. Gerhard Mann Chem.-Pharm. Fabrik, Berlin, Germany) was used to keep the eyes moist throughout the surgical procedure. After trimming the hair from the head of the animal, the surface of the scalp was disinfected with Betadine (Mundipharma, Cambridge, England) and 70% alcohol. To access the skull, a small incision was made in the skin along the midline, between the eyeline and the lambda, then four pean forceps were used to retract the skin from the top of the skull. After that, the connective tissue was removed from the skull, then a small amount of dental etching gel (Etch-Rite, PULPDENT Corporation, Watertown, MA, USA) containing 38% phosphoric acid was applied to the cleaned area to roughen the skull surface for better adhesion of the dental cement used to secure the 3D-printed fixture. For additional stabilization of the fixture, we mounted four bone screws (0.9 mm in diameter and 2 mm in length; ATLAS Neuroengineering, Leuven, Belgium) in the skull over the right hemisphere. A fifth bone screw (0.9 mm in diameter and 3 mm in length) with a wire soldered to it and serving as the reference and ground electrode for the recordings, was placed above the cerebellum. Using a dental drill, a 2 mm × 2 mm rectangular craniotomy was prepared over the thalamic area of interest (posterior and lateral posterior thalamic nuclei; AP: –3.6 mm, ML: 2.5 mm, with respect to the bregma). The dura mater was then carefully punctured above the insertion site with a sterile 34-gauge needle to reduce brain dimpling during probe insertion. To keep the neocortex hydrated until the probe was implanted, sterile room temperature physiological saline solution was regularly dripped into the cranial cavity. The Neuropixels probe attached to the 3D-printed fixture (mounted on a motorized stereotaxic micromanipulator) was positioned over the insertion site. Next, a mixture of bone wax and paraffin oil was applied to the base of the probe shank to create a hermetic seal to protect the probe from the dental cement. The probe was then implanted into the brain tissue to a depth of 7 mm from the cortical surface using a slow insertion speed (2 μm/s) to reduce the insertion-related mechanical tissue damage. Following probe implantation, we sealed the exposed cortex with a soft, two-component silicone gel (DuraGel, Cambridge NeuroTech, Cambridge, United Kingdom) to reduce the risk of inflammation and cerebrospinal fluid leakage. Next, the skull connector was secured to both the surface of the skull and the bone screws with light-curing dental cement (RelyX Universal Resin Cement, 3M, St. Paul, MN, USA). After that, the reference electrode was connected to the pin protruding from the skull connector, then the skin of the animal was sutured around the fixture with 2-3 stitches. Finally, while the animal was still under light anesthesia, a short (∼10 min) recording was acquired to inspect the signal quality and to record thalamic activity in the anesthetized state. After that, the rat was transferred to the animal facility for a five-day recovery period during which it received antibiotics (Augmentin, Glaxo Wellcome Production, Mayenne, France; 20 mg/kg body weight) and analgesics (Panadol Baby, Farmaclair, Hérouville-Saint-Clair, France; 2 mg/ml).

#### 2.3.2. Electrophysiological recording sessions

For the recording sessions, the animal was transferred from the animal facility to a dark and soundproof Faraday cage, then it was connected to the Neuropixels recording system. We started recording when the movement artifacts on the continuously monitored thalamic signals decreased significantly, indicating that the animal was probably starting its first sleep cycle. We recorded thalamic activity on a daily basis for about two weeks starting on the fifth day after probe implantation. The recording sessions varied in duration (0.5 – 3 hours) but a single recording session usually included several sleep-wake cycles. The parameters of data acquisition were the same as described above for the acute Neuropixels experiments.

### 2.4. Histology

To detect the tracks of silicon probes in the brain tissue and to identify thalamic nuclei, we applied a histological procedure described previously (Fiath et al., 2016b; Fiath et al., 2019). Briefly, after collecting the thalamic recordings, animals were deeply anesthetized and transcardially perfused with physiological saline solution (100 ml) followed by a fixative solution containing 4% paraformaldehyde in 0.1M phosphate buffer (pH = 7.4, 250 ml). The brain was removed from the skull after fixation and stored overnight at 4°C in the same fixative solution. Histological processing started by cutting 60-µm-thick coronal sections from the fixed brain with a vibratome (Leica VT1200, Leica Microsystems, Wetzlar, Germany). For rats, where the shanks of multi-shank probes were implanted along the anterior-posterior axis, sagittal sections were prepared. Following washing in 0.1 M phosphate buffer, brain sections were first placed in a Petri dish containing gelatin, then they were mounted onto microscopic slides and air dried. To identify brain sections containing fluorescent marks of DiI corresponding to probe tracks, the slides were examined under a light microscope (Leica DM2500, Leica Microsystems) equipped with a fluorescence LED illumination source (SFL4000, Leica Microsystems) and with a digital camera (DP73, Olympus, Tokyo, Japan). After that, the brain sections were processed for cresyl violet (Nissl) staining, dehydrated in xylene and coverslipped with DePex (SERVA Electrophoresis, Heidelberg, Germany). Finally, Nissl-stained sections containing the tracks of silicon probes were photographed under the microscope.

### 2.5. Data analysis

#### 2.5.1. Detection of up-state onsets and computation of the up-state onset locked multiunit activity average

In recordings obtained from anesthetized animals, we used a multiunit activity (MUA)-based method to identify the onset of thalamic (or cortical) up-states, a modified version of the procedure described in our previous studies (Fiath et al., 2016a; Fiath et al., 2016b). The detection process was similar for both multi-shank and single-shank (Neuropixels) recordings. First, we removed low frequency components (e.g., local field potentials for wideband recordings obtained with multi-shank probes) from the recordings using a bandpass filter (third-order Butterworth bandpass filter, 500 – 5000 Hz, zero-phase shift) and rectified the filtered signal (resulting in the MUA signal). Then, the MUA was downsampled by a factor of ten for faster processing (e.g., from 20 kHz to 2 kHz for multi-shank recordings). After that, the samples of the downsampled MUA were summed up across all channels resulting in a single channel time series representing the instantaneous intensity of population activity in the thalamus (or the cortex). For Neuropixels recordings, before summing up the samples, we removed all channels located outside the thalamus (e.g., hippocampal channels). Next, the summed signal was smoothed with an additional low-pass filter (third-order Butterworth filter, 50 Hz, zero-phase shift) to obtain the smoothed population activity (SPA). In most of the cases, the amplitude histogram of the SPA showed a clear bimodal distribution.

To determine the onset of up- and down-states, we first calculated an amplitude threshold level. This was done by estimating the average (AVG) and standard deviation (SD) of the thalamic/cortical activity measured during down-states (lack of action potential firing) based on the SPA signal by identifying the center of down-states and calculating the average amplitude within a 50 ms long window around this center. Based on empirical observations, a threshold level of AVG + (2.5 * SD) was usually sufficient to accurately determine the onset of up- and down-states. The state detection algorithm was configured with a minimum duration of 50 ms for up-states and a minimum duration of 100 ms for down-states. For recordings where the onset detection was less accurate, the value of the multiplier of the SD and the minimum state durations were fine-tuned. The onset of an up-state was defined as the time point when the amplitude value of the SPA signal exceeded the calculated threshold level. This time point had to be preceded by a down-state with a duration equal to or longer than the defined minimum state duration. The extracted up-state onset times were saved to files. Finally, to reduce the effect of outliers (i.e., very short or very long up-states which are often the result of incorrect or inaccurate state detections) on the up-state onset locked thalamic/cortical MUA averages, we kept only up-states with a duration of AVG ± (1.5 * SD). To compute these MUA averages (termed as MUA depth profiles for thalamic recordings), short snippets were cut from the continuous MUA recordings (filtered between 500 – 5000 Hz and rectified) around the up-state onset (100/150 ms before and 300 ms after the onset) and averaged across up-states. Finally, the MUA average was smoothed using a third-order Butterworth filter (50 Hz, low-pass, zero-phase shift), then z-score normalization was applied to the smoothed MUA average.

#### 2.5.2. Identification of preferred propagation patterns during thalamic up-states in Neuropixels recordings

To identify preferred directions of activity propagation in the thalamus (e.g., dorsal-to-ventral or ventral-to-dorsal), we selected several Neuropixels recordings that exhibited clear MUA propagation (i.e., propagation was evident also in the up-state onset locked MUA depth profiles; n = 15 recordings from 14 rats), then identified the approximate dorsal and ventral thalamic locations (channels) where propagation started and terminated in these recordings (i.e., the borders of propagation). Next, we selected ten adjacent channels with at least moderate MUA levels at three thalamic positions: at the previously determined ventral and dorsal propagation borders, and midway between the two borders. After that, we identified the onset of up-states separately at these three thalamic locations using the state detection algorithm described above. Only up-states that were detected at all three locations within a short time frame were kept for further analysis (i.e., within a time window of ±200 ms with respect to up-state onsets detected at the ventral thalamic location used as reference). Delays of up-state onsets between the three thalamic locations were calculated, then the sequence of MUA initiation (propagation pattern) was determined for each up-state. Finally, the rate of occurrence and the proportion of all six possible propagation patterns were calculated.

#### 2.5.3. Correlation between cortical and thalamic propagation patterns during up-states

For the experiments with simultaneous cortical and thalamic recordings (n = 5 recordings from 5 rats), to determine the temporal relationship between cortical and thalamic slow waves, MUA averages were calculated in both brain areas using either the onset of cortical up-states (detected based on the SPA of all cortical channels), or the onset of thalamic up-states (detected based on the SPA of all thalamic channels or based only on thalamic channels displaying MUA propagation), in the same way as described above. In addition, to investigate the correlation between cortical and thalamic propagation patterns, we detected up-state onsets at three cortical and three thalamic positions. For the cortex, these positions were the following: between shanks 1 and 2 (by detecting up-state onsets on both shanks and calculating their average; anterior position), between shanks 4 and 5 (middle position) and between shanks 7 and 8 (posterior position). Based on the up-state onset delays between cortical positions, we defined the direction of cortical activity propagation during up-states (e.g., anterior-to-posterior or posterior-to-anterior). Thalamic propagation patterns were determined in the same manner as described above. Finally, the rate of occurrence and the proportion of all six possible propagation patterns were calculated both for the cortex and thalamus.

#### 2.5.4. Sleep stage scoring in chronic recordings

To identify periods of slow-wave sleep and to detect activity propagation in the thalamus of chronically implanted animals, we first computed spectrograms from local field potential recordings in the frequency range between 0.1 and 100 Hz on a channel located in the dentate gyrus region of the hippocampus (Supplementary Fig. S1). Spectrograms were calculated using Chronux (v2.12; Bokil et al., 2010). Next, we computed the mean power in the theta (5.5 - 8 Hz) and the delta (1 - 4 Hz) frequency bands, as well as the ratio of the mean power in these two frequency bands (theta/delta ratio; Supplementary Fig. S1). These spectral features allowed us to reliably separate non-rapid eye movement (NREM) sleep from rapid eye movement (REM) sleep and wake states (Rayan et al., 2022). REM sleep and wake states were characterized by an increased theta power and a decreased delta power (high theta/delta ratio), whereas NREM sleep was marked by decreased theta and increased delta power (low theta/delta ratio; Supplementary Fig. S1). Furthermore, REM and wake states could be distinguished by a difference in the mean power of the high gamma band (70 - 100 Hz): REM sleep was characterized by an enhanced high gamma activity compared to wakefulness (Supplementary Fig. S1). Additionally, a weaker theta rhythm and movement artifacts were further indications of wake states. These observations (increased high gamma and theta power during REM sleep compared to wake) are also in line with the findings of Montgomery et al. (2008). Thalamic activity during the identified NREM sleep periods was then visually inspected to detect and mark MUA events exhibiting propagation. In general, a single NREM sleep period with a duration of at least 5 minutes was examined for each recording session (average duration of examined NREM periods: 12.04 ± 4.66 min). In total, 21 recording sessions were analyzed for the two chronically implanted rats (13 and 8 sessions, respectively). Finally, we calculated the rate of occurrence of the identified propagating MUA events, and computed MUA depth profile averages locked to the onset of these events.

#### 2.5.5. Identification of thalamic nuclei

Thalamic nuclei from which we recorded population activity were identified on the basis of the Nissl-stained brain sections, the verified position of the probe tracks and neuronal activity in electrophysiological recordings. In some cases, sensory stimulation (somatosensory, auditory or visual) was performed to aid the identification of the location of sensory (relay) thalamic nuclei. To demarcate thalamic nuclei in Nissl-stained thalamic sections containing the probe tracks, we first identified major anatomical landmarks in these sections (e.g., parts of the hippocampus, internal capsule) then used the sections in the stereotaxic atlas of the rat/mouse brain for comparison (Paxinos and Franklin, 2004; Paxinos and Watson, 2007). The results were fine-tuned using information obtained from spontaneous and stimulation-evoked neuronal recordings. In the case where a notable discrepancy between the original implantation coordinates and the actual anatomical position of the probe was detected based on the obtained anatomical and electrophysiological information (>200 µm difference in AP, ML or DV coordinates), we updated the implantation coordinates to improve the accuracy of the colormaps summarizing the location of thalamic recordings in rats (Fig. 2 and Supplementary Fig. S5). Brain regions and their abbreviations used in the text and figures are listed in Supplementary Table S1.

**Figure 1.**
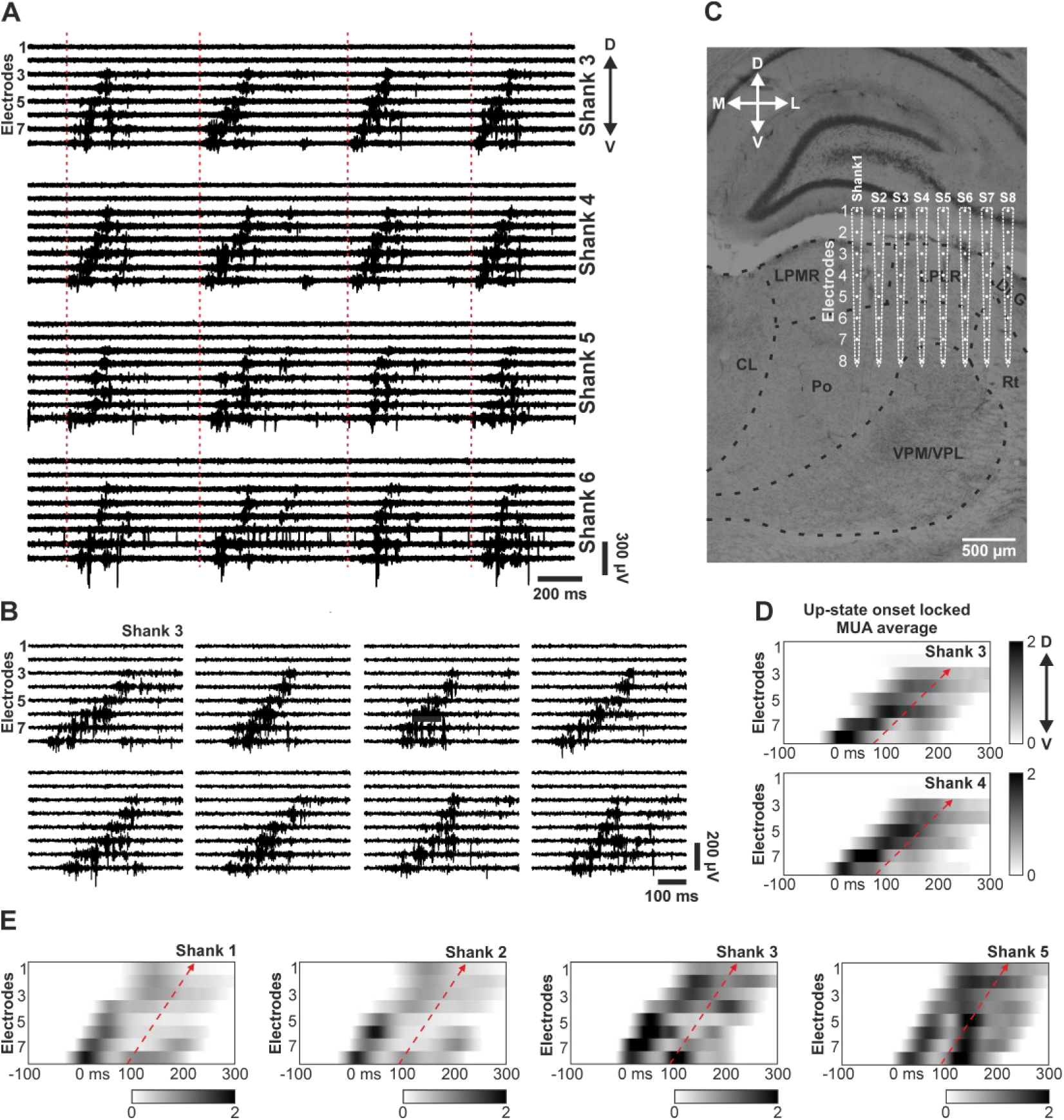
Propagating thalamic population activity observed with multi-shank silicon probes under ketamine/xylazine anesthesia in rats. (A) Representative spontaneous thalamic multiunit activity (MUA) traces recorded on four adjacent shanks in the posterior (Po) and lateral posterior (LP) thalamic nuclei. Up-state onsets are indicated by vertical dashed red lines. Activity propagation from ventral (bottom electrodes) to dorsal (top electrodes) regions of the thalamus is visible on shanks 3 and 4. (B) Examples of individual thalamic up-states exhibiting propagating MUA recorded on shank 3 at various time points of the 30-minute-long recording. (C) Nissl-stained coronal brain section illustrating the approximate location of the shanks and recording sites (electrodes) of the silicon probe relative to major thalamic nuclei. Shanks 3 and 4 recorded mainly from two higher order nuclei (Po and LP). (D) Normalized (z-score) MUA depth profile averages locked to the up-state onsets (n = 2452 up-states) on shanks 3 (top) and 4 (bottom) showing ventral-to-dorsal propagation of thalamic activity (dashed red arrow). The up-state starts at time point zero. (E) Normalized (z-score) up-state onset (n = 2857 up-states) locked MUA depth profile averages on four shanks located in the dorsal thalamus (Po and LP) of another rat. D – dorsal; V – ventral; M – medial; L – lateral.

**Figure 2.**
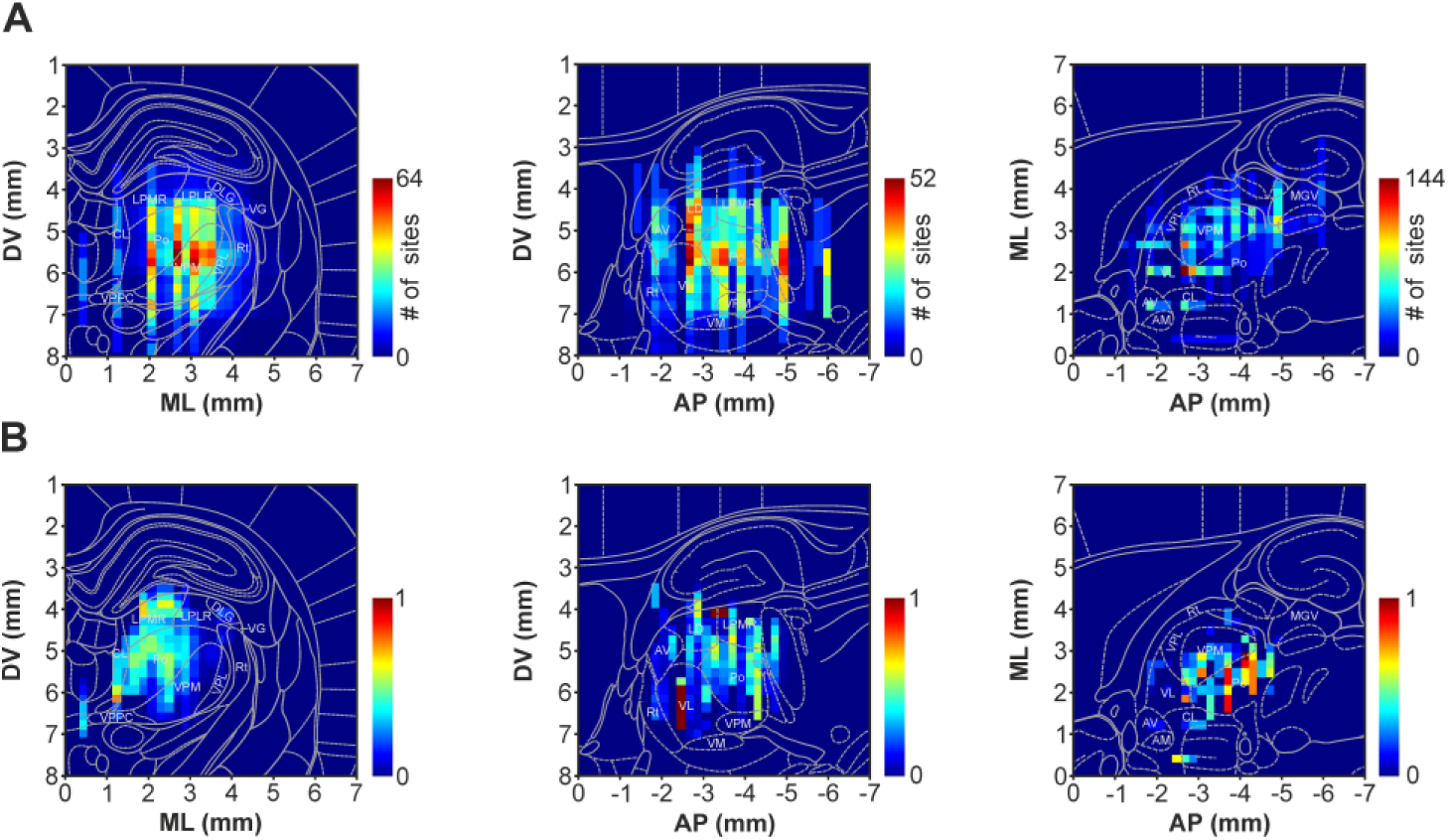
Colormaps illustrating the distribution of recording site locations used to collect thalamic activity with multi-shank silicon probes in anesthetized rats (A) along the three anatomical axes (left: anterior-posterior (AP) axis; middle: medial-lateral (ML) axis; right: dorsal-ventral (DV) axis), and the probability of observing propagating activity along the dorsoventral axis at these sites (B). Each pixel within these colormaps represents a small quadratic region in the brain with an area of 0.2 mm x 0.2 mm. The pixel values represent cumulative values along one of the three anatomical axes. Schematic brain sections overlaid on the colormaps indicate the location of relevant thalamic nuclei at coordinates AP -3.6 mm (left), ML 1.9 mm (middle) and DV 5.6 mm (right), relative to the bregma.

## 3. Results

### 3.1. Propagation of neuronal population activity is common during spontaneous up-states in dorsal thalamic nuclei of rats under anesthesia

To investigate whether propagation of population activity occurs within the thalamus of rats, we first implanted multi-shank silicon probes to get a qualitative picture of the spatiotemporal dynamics of spontaneous thalamic activity patterns during ketamine/xylazine-induced slow waves. Ketamine/xylazine anesthesia in rats induces slow oscillation in the thalamocortical network with a peak frequency of approximately 1.5 Hz and with a regular and stable pattern of alternating up- and down-states (David et al., 2013; Fiath et al., 2016b; Sharma et al., 2010). In 14 rats, we performed a total of 40 thalamic insertions with an eight-shank, 64-electrode silicon probe. In these recordings, we consistently observed clear multiunit activity (MUA, 500 – 5000 Hz) propagation during up-states in various thalamic nuclei (Figs. 1 and 2). Our definition of MUA propagation was a clearly visible delay (> 10 ms) in the onset of up-states on at least three adjacent channels/electrodes (the distance between adjacent electrodes and shanks of multi-shank probes was 200 µm), which delay increased gradually as a function of distance from the channel where the spiking activity initiated (i.e., the channel with the earliest up-state onset). These propagating up-states could typically be recorded simultaneously on multiple electrodes (3-8) and on multiple probe shanks (2-5), with a propagation distance spanning several hundred micrometers in all directions (Fig. 1A and B; Supplementary Figs. S2 and S3). The time lag between up-state onsets at the starting channel and the channel with the longest onset delay could extend to several hundred milliseconds (∼100 ms on average; Fig. 1D and E; Supplementary Figs. S2 and S3). Based on both visual inspections of thalamic recordings and the computed up-state onset locked MUA depth profile averages, alongside with the evaluation of histological results (e.g., Fig. 1C), propagating population activity was most frequently observed in the dorsal thalamus, more specifically in higher order nuclei such as the posterior (Po), the lateral posterior (LP) or the laterodorsal (LD) nucleus (Fig. 2). In contrast, thalamic activity propagation was less common or virtually absent in ventral thalamic structures (except for propagation of sleep spindles; see more details below) such as the ventral posteromedial nucleus (VPM; Fig. 3A) or the ventral division of the medial geniculate nucleus (MGV; Fig. 3B). Within these particular thalamic nuclei, up-states started quasi-synchronously on adjacent channels (Fig. 3A and B).

**Figure 3.**
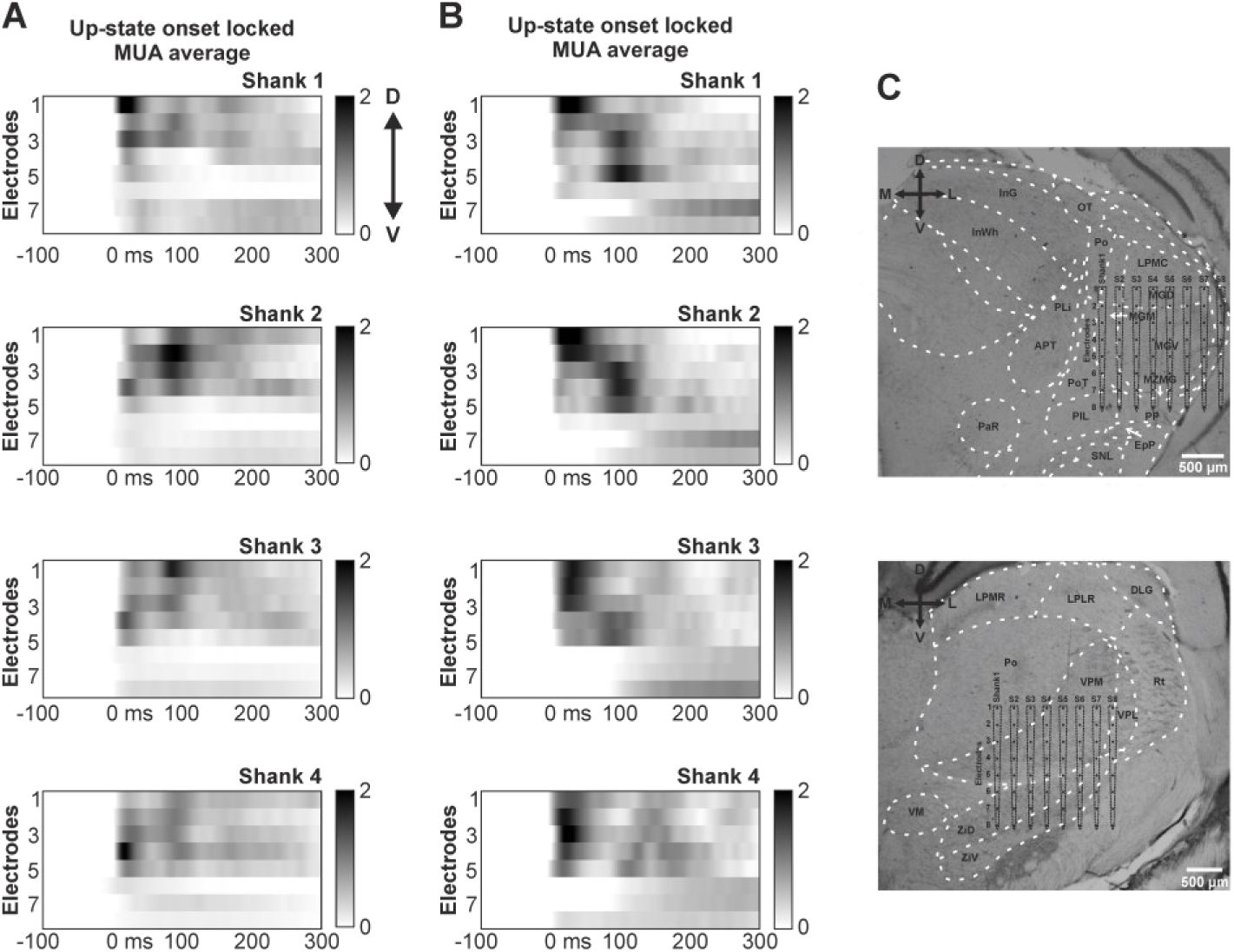
Synchronous multiunit activity at the up-state onset observed in the thalamus with multi-shank probes in anesthetized rats. (A-B) Normalized (z-score) up-state onset locked MUA depth profile averages on four shanks located in ventral (A; ventrobasal complex; n = 1285 up-states) and posterior (B; auditory thalamic nuclei; n = 2269 up-states) regions of the thalamus. The up-state starts at time point zero. Note that MUA at the up-state onset starts nearly simultaneously on different electrodes. (C) Nissl-stained coronal brain sections illustrating the approximate location of the shanks and electrodes of the silicon probe relative to major thalamic nuclei (top brain section corresponds to panel A; bottom brain section corresponds to panel B). D – dorsal; V – ventral; M – medial; L – lateral.

In thalamic areas where up-state propagation was observed (n = 43/82 recordings in 14 rats; ∼52%), the direction of propagation was predominantly along the dorsal-ventral axis (usually from ventral to dorsal; n = 41/82 recordings; ∼50%). This propagation trend was also commonly visible in the up-state onset locked MUA depth profiles, suggesting a consistent and stereotypical pattern of propagation with minimal variability (Fig. 1). While alternate propagation patterns, such as dorsal-to-ventral propagation of neuronal activity, could also be observed, their rate of occurrence was comparatively lower. We also investigated propagation patterns along the anterior-posterior or the medial-lateral axis (across probe shanks). Although we could detect posterior-to-anterior propagation (n = 5/27 recordings in 2 out of 4 rats; ∼19%; Supplementary Fig. S2) as well as medial-to-lateral propagation (n = 5/55 recordings in 4 out of 10 rats, ∼9%; Supplementary Fig. S3) of thalamic MUA activity, these propagation patterns were less prevalent than activity propagation along the dorsoventral axis (∼50% vs. ∼9%/∼19% of recordings, respectively). These posterior-to-anterior and medial-to-lateral propagation patterns were also less consistent and regular, and tended to have shorter propagation delays (< 100 ms). Thus, in the rest of the study we focused on the characterization of thalamic up-states propagating along the dorsoventral axis.

Compared to single-shank devices, the application of neural probes with multiple silicon shanks may lead to a larger brain dimpling during implantation, along with more severe insertion-related mechanical tissue damage. This can be attributed to the substantially larger total volume occupied by the implanted shanks, which results in an increased displacement and deformation of the brain tissue (Chen et al., 2021; Szarowski et al., 2003). For instance, during acute experiments, we frequently noticed bleeding at the cortical surface, or traces of blood around the track of several probe shanks in brain sections after histological processing (Supplementary Fig. S4A). In some of these experiments, as well as in a few other cases, we observed very low amplitude or absent MUA (i.e., lack of spiking activity) on certain shanks which may be an indication of severe tissue damage (Supplementary Fig. S4B). Given that damage to the brain tissue (particularly in proximity to the recording sites) might significantly affect the ongoing neuronal activity, the use of a less invasive neural probe could provide a clearer and more reliable picture on the spatiotemporal details of thalamic MUA propagation. Therefore, after we identified potential thalamic areas displaying propagation of population activity by using multi-shank probes, we proceeded to examine thalamic slow waves with single-shank, high-density Neuropixels silicon probes in 20 anesthetized animals (n = 46 insertions; Table 1; Supplementary Fig. S5), focusing mainly on these specific areas. Furthermore, in addition to the reduced shank cross-section of this neural probe type, which mitigates the insertion-related tissue damage, they also provide an enhanced spatial sampling of neuronal activity due to their high electrode density. For instance, in our recordings, generally about 300 electrodes collected the activity of multiple thalamic nuclei simultaneously.

Spontaneous thalamic activity obtained with Neuropixels probes from anesthetized rats typically showed a gradual propagation of MUA within and between dorsal thalamic nuclei such as the Po (Fig. 4A), the LP (Fig. 4A and B) or the dorsal lateral geniculate nucleus (DLG; Fig. 4C; propagation was detected in 34 out of 41 recordings, ∼83%). While the most prominent propagation pattern observed was the ventral-to-dorsal propagation of MUA, also evident in the up-state onset locked MUA depth profiles, other patterns could also be detected (e.g., dorsal-to-ventral propagation, activity originating at a specific point of the thalamus and subsequently traveling both dorsally and ventrally, or synchronous MUA; Supplementary Fig. S6). In contrast, MUA in ventral parts of the thalamus initiated nearly simultaneously at the up-state onset (Fig. 4A-C). In other thalamic areas, such as the ventral or the dorsal parts of the medial geniculate nucleus (MGV and MGD, respectively), up-states also started relatively synchronously at distinct thalamic depths (Fig. 4D).

**Figure 4.**
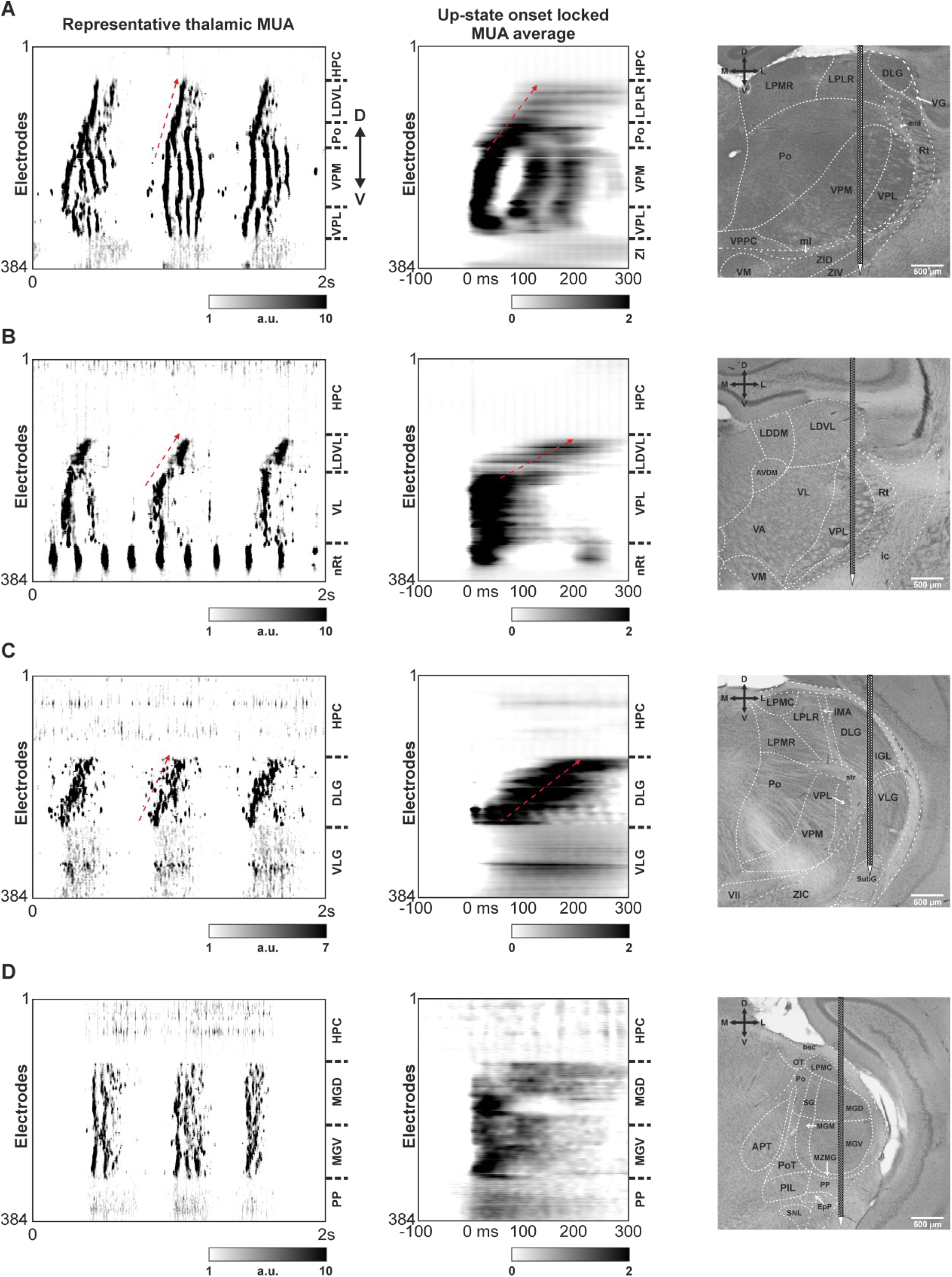
Propagating thalamic population activity observed with single-shank high-density silicon probes under ketamine/xylazine anesthesia in rats. (A-C) Left: two-second-long recordings of spontaneous multiunit activity from different thalamic regions demonstrating the characteristic ventral-to-dorsal pattern of MUA propagation during up-states. To improve the visualization of neuronal activity, the grayscale MUA maps are smoothed both in time (50 Hz low-pass filter) and in space (data from two adjacent electrodes are averaged). Boundaries between major thalamic nuclei are marked by horizontal dashed lines on the right side of the MUA maps. Oblique dashed red arrows indicate the direction of propagation. Middle: normalized (z-score) up-state onset locked MUA depth profile averages (n = 2398, 3488 and 1787 up-states for panels A-C, respectively). The up-state starts at time point zero. Right: Nissl-stained coronal brain sections illustrating the approximate location of the probe shank and electrodes relative to major thalamic nuclei. (D) Another thalamic recording position without visible MUA propagation (the MUA average was computed from n = 645 up-states). D – dorsal; V – ventral; M – medial; L - lateral.

It is important to note that MUA during thalamic sleep spindles (e.g., in the VPM/VPL, see Fig. 4A) also exhibited propagating behavior, however, this form of propagation differed from MUA propagation observed during up-states: it was faster (indicated by a steeper activity propagation during spindle cycles), less consistent and more variable (more spindle cycles are included in a single up-state and each spindle cycle could show a distinct propagation pattern). Furthermore, we can see in the thalamic MUA depth profile that, on average, sleep spindle-related MUA in the VPM/VPL nuclei started nearly synchronously during the up-state onset. Interestingly, however, a less obvious propagation pattern can be discerned in consecutive spindle cycles (Fig. 4A). Since our focus was on characterizing the propagation of slow waves, we did not analyze propagation patterns during sleep spindles further.

In summary, thalamic MUA during up-states shows propagating behavior in ketamine/xylazine anesthetized rats. However, propagation is mostly confined to the dorsal parts of the thalamus, where mainly higher order nuclei are located. The distance of propagation ranged from several hundred micrometers to up to 1.8 millimeter (1.14 ± 0.39 mm), while the propagation delay was approximately 100 ms, resulting in a propagation speed of 14.6 ± 8.6 mm/s (measured in n = 13 Neuropixels recordings).

### 3.2. Dorsal thalamic up-states propagate most frequently in the ventral to dorsal direction

Based on visual assessment of thalamic recordings from anesthetized rats, it is evident that the dominant direction of thalamic propagation along the dorsoventral axis is from ventral to dorsal (see Figs. 1, 4 and 5A). However, other propagation patterns could also be observed, such as propagation in the opposite direction (Fig. 5B). The speed of these propagation patterns was variable, it could be faster or slower (Fig.5A and B, left vs. right) but the propagation had an average speed of about 15 mm/s. Next, by selecting 15 recordings with clear and consistent thalamic propagation activity, we quantified the rate of occurrence and the proportion of distinct propagation patterns. To perform this, we first defined three equidistant locations in the region of the thalamus exhibiting MUA propagation: a dorsal location close to the dorsal border of activity propagation, a ventral point next to the ventral border of activity propagation and a location halfway between the two borders (see the Methods section for more details; Fig. 5C). Then, for each up-state, we determined the sequence of MUA initiation at these positions and calculated the frequency of different sequences.

**Figure 5.**
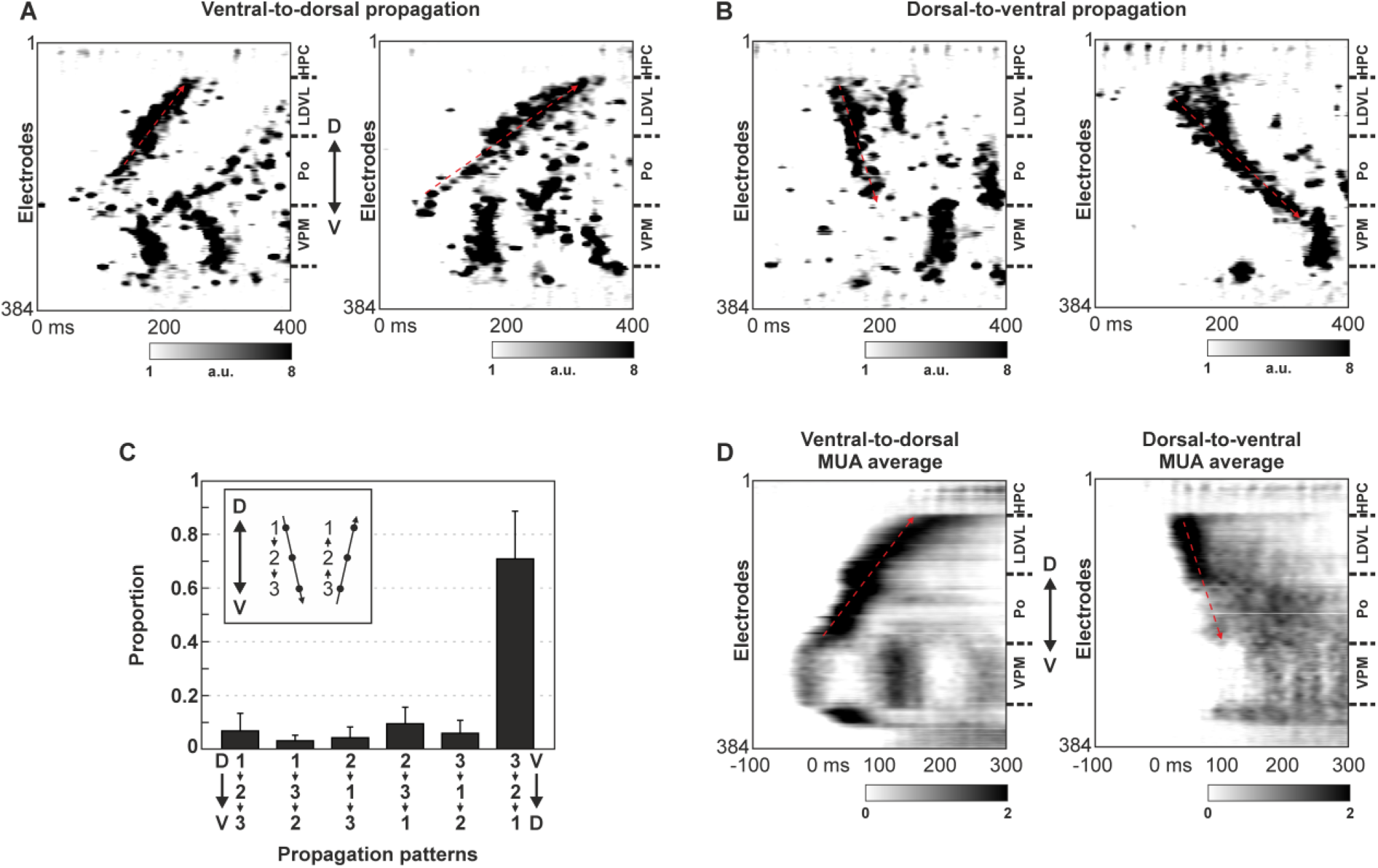
Distinct propagation patterns can be observed in the dorsal thalamus of anesthetized rats, but propagation in the ventral-to-dorsal direction is the most frequent. (A) Two examples of the characteristic ventral-to-dorsal MUA propagation in the dorsal thalamus. Note the difference in propagation speed between the two up-states. (B) Two examples of the less frequent dorsal-to-ventral propagation pattern recorded at the same thalamic location. Note the difference in the propagation speed between the two up-states. (C) Proportion of different thalamic propagation patterns (n = 15 recordings from 14 rats). Sequence 1-2-3 corresponds to dorsal-to-ventral propagation, while sequence 3-2-1 indicates the proportion of up-states with ventral-to-dorsal propagation. Thalamic locations: 1 – most dorsal, 2 – midway between locations 1 and 3; 3 – most ventral. (D) Normalized (z-score) up-state onset locked MUA depth profile averages for the ventral-to-dorsal (left, n = 980 up-states) and dorsal-to-ventral (right, n = 335 up-states) propagation patterns in a single animal. MUA averages were calculated from the same recording from which examples are shown in panels A and B. Boundaries of major thalamic nuclei are marked by horizontal dashed lines on the right side of the MUA maps. Oblique dashed red arrows indicate the direction of propagation. The up-state starts at time point zero. D – dorsal, V – ventral.

Among the six possible patterns, the ventral-to-dorsal propagation was by far the most frequent with approximately two out of three up-states displaying this particular pattern (∼71%; Fig. 5C). The remaining five patterns occurred at much lower frequencies, ranging from 3-10% (Fig. 5C). The presence of distinct propagation patterns is further supported by the up-state onset locked MUA depth profiles calculated using up-states corresponding to these individual patterns (e.g., ventral-to-dorsal and dorsal-to-ventral propagation patterns; Fig. 5D). Consequently, we can conclude, that although the ventral-to-dorsal propagation pattern is by far the most common pattern observed, other propagation patterns (including more complex patterns; Supplementary Fig. S6) are also present in the rat thalamus under anesthesia.

### 3.3. Spontaneously occurring dorsal thalamic up-states are temporally locked to cortical up-state activity

Thalamic nuclei establish complex, reciprocal connections with numerous cortical areas. Furthermore, previous studies have found that spontaneous slow waves in the neocortex of rodents propagate across cortical areas, predominantly along the anterior-posterior axis (Greenberg et al., 2018; Greenberg and Dickson, 2013; Ruiz-Mejias et al., 2011; Stroh et al., 2013). After exploring the spatiotemporal dynamics of slow waves in the thalamus, we became interested in the temporal relationship between cortical and thalamic MUA during up-states. Therefore, in a subset of experiments (n = 5 rats), in addition to the Neuropixels probe inserted into the thalamus (recording from Po, LD and VPM nuclei), we implanted an eight-shank silicon probe into the neocortex with its shanks aligned along the anterior-posterior axis. In these simultaneous thalamocortical recordings, we found that MUA during spontaneous cortical up-states usually started at the electrodes of anteriorly located shanks (or quasi-synchronously on all shanks), while thalamic activity showed the characteristic ventral-to-dorsal propagation pattern with MUA starting at the ventral part of the Po and terminating at the dorsal part of the LD nucleus (Fig. 6). To examine the possible influence of the neocortex on thalamic activity patterns, we calculated and compared MUA depth profiles locked to either cortical (Fig. 6 B and D) or thalamic up-state onsets (either including all thalamic channels for up-state onset detection, see Supplementary Fig. S7A; or including exclusively those dorsal thalamic channels that displayed activity propagating, see Supplementary Fig. S7B). Interestingly, thalamic MUA depth profiles where the ventral-to-dorsal propagation was most evident were those locked to cortical up-state onsets (compare Fig. 6B and D vs. Supplementary Fig. S7). This suggests that MUA in the dorsal thalamus is tightly locked in time to cortical up-states, and that the neocortex may have a noticeable influence on ongoing thalamic propagation patterns. Another interesting observation was that in 4 out of 5 rats, up-state-related MUA started first in the ventral part of the Po, leading the onset of the cortical up-state by about 50 ms (Fig. 6B and D). Propagation speed of thalamic MUA was measured 12.4 ± 4.5 mm/s, whereas cortical MUA propagated about three times faster (37.9 ± 18.0 mm/s).

**Figure 6.**
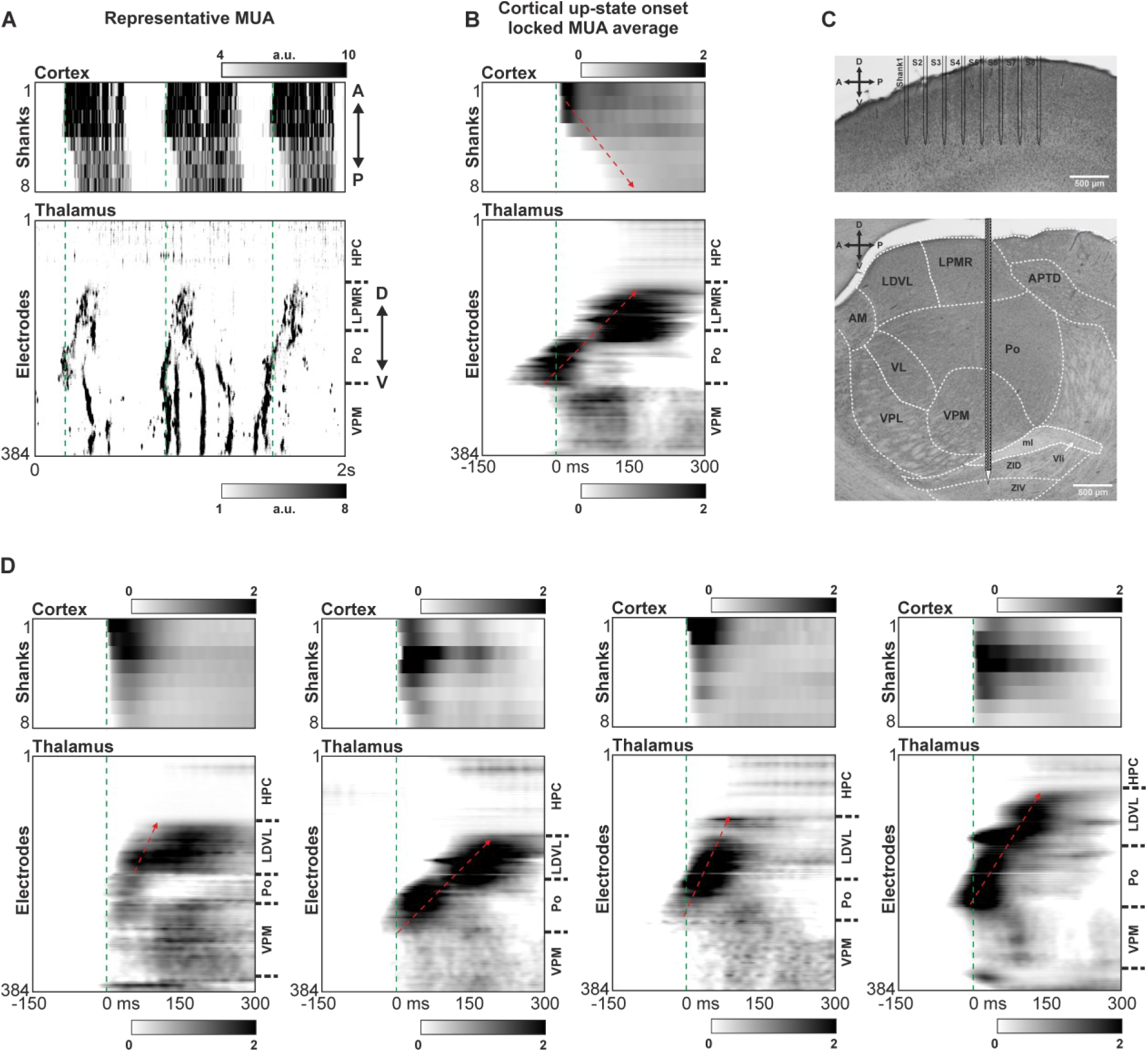
Simultaneous cortical and thalamic multiunit activity recordings in anesthetized rats display propagation of neuronal activity during up-states in both regions. (A) Two-second-long representative recordings from the neocortex (top, multi-shank probe) and the thalamus (bottom, Neuropixels probe) demonstrating anterior-to-posterior propagation in the neocortex and the characteristic ventral-to-dorsal propagation in the thalamus. For the neocortex, MUA recorded by all electrodes (n = 8) located on a single shank was averaged for each shank. (B) Normalized (z-score) cortical up-state onset locked cortical (top) and thalamic (bottom) MUA depth profile averages (n = 3420 up-states) of one of the experiments. (C) Nissl-stained sagittal brain section illustrating the approximate location of the shanks and electrodes of the two silicon probes in the neocortex (top) and the thalamus (bottom). (D) Cortical up-state onset locked MUA depth profile averages in the other four rats (n = 4159, 1803, 2341 and 5141 up-states from left to right, respectively). Boundaries of major thalamic nuclei are marked by horizontal dashed lines on the right side of the MUA maps. Oblique dashed red arrows indicate the direction of propagation. Vertical dashed green lines mark the cortical up-state onsets. A – anterior, P – posterior, D – dorsal, V – ventral.

To examine whether there is a correlation between cortical and thalamic propagation patterns, we first determined the proportion of the main cortical and thalamic MUA patterns, similarly to that described above for the thalamus. In the neocortex, the dominant propagation pattern was the anterior-to-posterior traveling of up-states (∼49% of up-states, Fig. 7A), with every second up-state propagating in this direction; whereas in the thalamus, the most frequent pattern was the previously observed ventral-to-dorsal propagation, with a rate of occurrence matching that of anterior-to-posterior traveling cortical up-states (∼55%, Fig. 7B). Consequently, cortical anterior-to-posterior propagation patterns were frequently accompanied by ventral-to-dorsal MUA propagation in the thalamus, however, only with an incidence similar to the frequency of ventral-to-dorsal propagation in the thalamus (∼52% of up-states, Fig. 7C). The other thalamic patterns could also be observed during anterior-to-posterior cortical propagation with a similar rate of occurrence as found in the thalamus. Nevertheless, thalamic ventral-to-dorsal propagation was even more discernible in the MUA depth profiles locked to these anterior-to-posterior cortical up-state onsets (e.g., compare the third column of Fig. 6D with Fig. 7C). Additionally, propagation of cortical up-states in the opposite direction (posterior-to-anterior), which had a significantly lower occurrence (∼13% of up-states), was often observed in association with thalamic MUA propagation in the dorsal-to-ventral direction (opposite to the most frequent thalamic propagation direction, with ∼11% occurrence, Fig. 7D). Interestingly, in one of the examined rats, this relationship was evident even in the MUA depth profile locked to cortical up-states propagating in the posterior-to-anterior direction (Fig. 7D). Thus, although each thalamic propagation pattern can emerge in the thalamus during each of the cortical traveling patterns, our findings suggest that there may be a weak correlation between ongoing cortical and thalamic MUA patterns.

**Figure 7.**
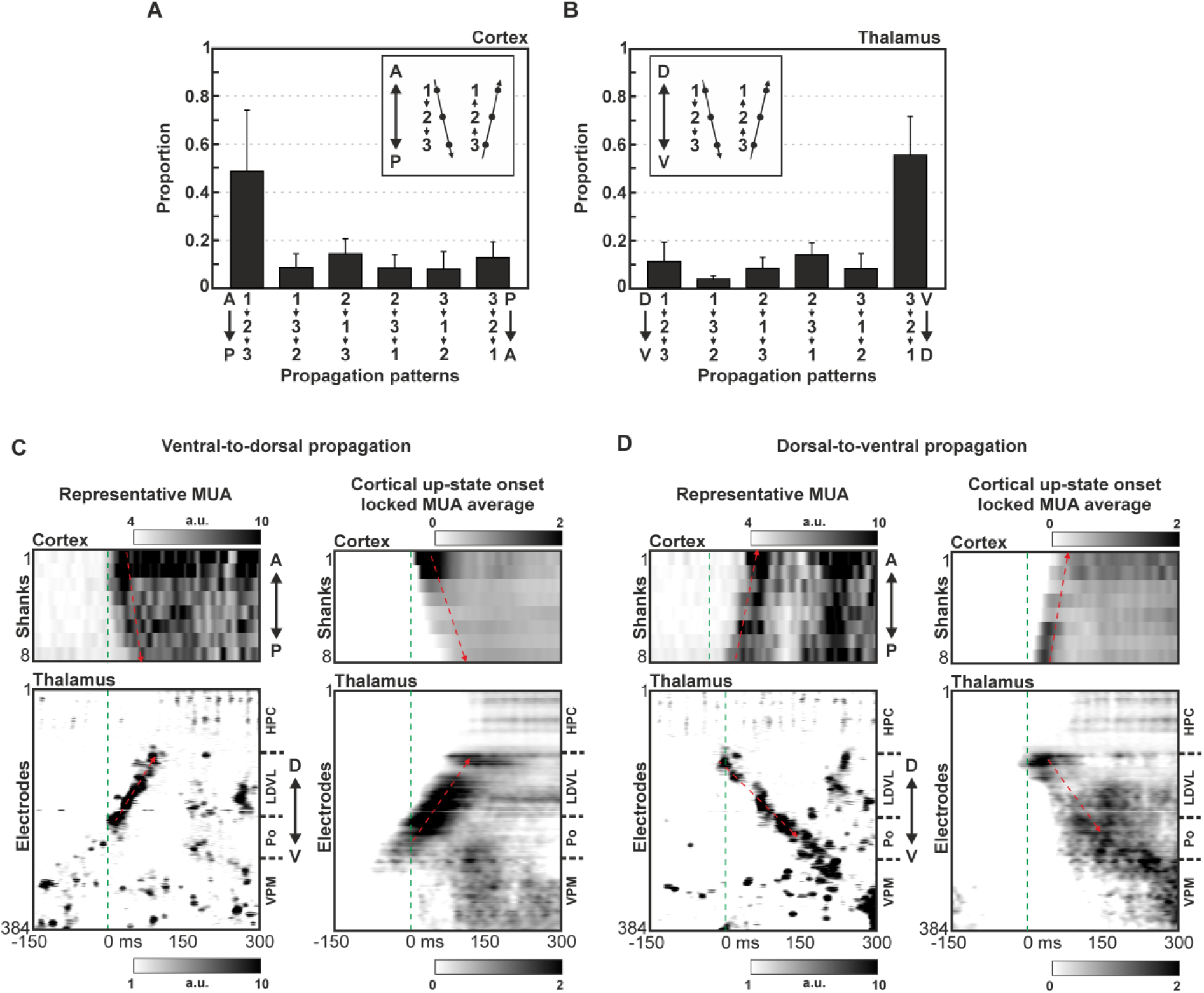
Cortical and thalamic propagation patterns observed during up-states are weakly correlated. (A) The proportion of distinct cortical propagation patterns (n = 5 thalamocortical recordings). Sequence 1-2-3 corresponds to anterior-to-posterior propagation, while sequence 3-2-1 indicates the proportion of up-states with posterior-to-anterior propagation. Cortical locations used to examine propagation patterns: 1 – between anteriorly located shanks 1 and 2, 2 – between shanks 3 and 4; 3 – between posteriorly located shanks 7 and 8. (B) The proportion of different thalamic propagation patterns in the same recordings. Sequence 1-2-3 corresponds to dorsal-to-ventral propagation, while sequence 3-2-1 indicates the proportion of up-states with ventral-to-dorsal propagation. Thalamic locations: 1 – most dorsally located electrodes in the thalamus displaying MUA propagation, 2 – midway between location 1 and 3; 3 – most ventral electrodes in the thalamus displaying MUA propagation. (C) Left: Example of a thalamic ventral-to-dorsal MUA propagation during a cortical up-state traveling in the anterior-to-posterior direction. Right: Normalized (z-score) cortical up-state onset locked cortical (top) and thalamic (bottom) MUA depth profile averages (n = 887 anterior-to-posterior cortical up-states). (D) Left: Example of a thalamic dorsal-to-ventral MUA propagation during a cortical up-state traveling in the posterior-to-anterior direction. Right: Normalized (z-score) cortical up-state onset locked cortical (top) and thalamic (bottom) MUA averages (n = 190 posterior-to-anterior cortical up-states). Boundaries of major thalamic nuclei are marked by horizontal dashed lines on the right side of the MUA maps. Oblique dashed red arrows indicate the direction of propagation. Vertical dashed green lines mark the cortical up-state onsets. A – anterior, P – posterior, D – dorsal, V – ventral.

### 3.4. Thalamic multiunit activity has similar propagation properties during spontaneous up-states in anesthetized mice

To investigate whether spontaneously occurring thalamic up-states display similar propagating behavior in other species, using Neuropixels (Fig. 8A and B) and multi-shank silicon probes (Fig. 8C and D), we recorded population activity from the thalamus of ketamine/xylazine-anesthetized mice. Although we were able to reliably observe the propagation of thalamic MUA in up-states, since the thalamus of mice is significantly smaller than the rat thalamus, the propagation of thalamic population activity was limited to smaller regions. Furthermore, because of this difference in the size of the thalamus, there was more uncertainty in the correct identification of the thalamic nuclei from which the activity was collected, making the demarcation of these nuclei more challenging. In mice, similarly to rats, thalamic MUA propagation was most frequently observed in the dorsal thalamus (in higher order nuclei such as the Po or LD) with the same dominant propagation pattern starting from ventral parts of the thalamus and traveling dorsally (Fig. 8). Like found in rats, other MUA patterns also occurred albeit with a lower frequency. Interestingly, although cortical up-state onset locked thalamic MUA depth profiles showed a discernible ventral-to-dorsal propagation in the examined mice, certain parts of the dorsal thalamus were not involved in the observed MUA propagation (e.g., see Fig. 8C). The propagation delay between the starting point of the up-state to its termination was also similar to that found in rats; namely, in the range of 100 to 300 ms, which, given the smaller size of the mouse thalamus, suggests a slightly slower propagation speed (< 5 mm/s).

**Figure 8.**
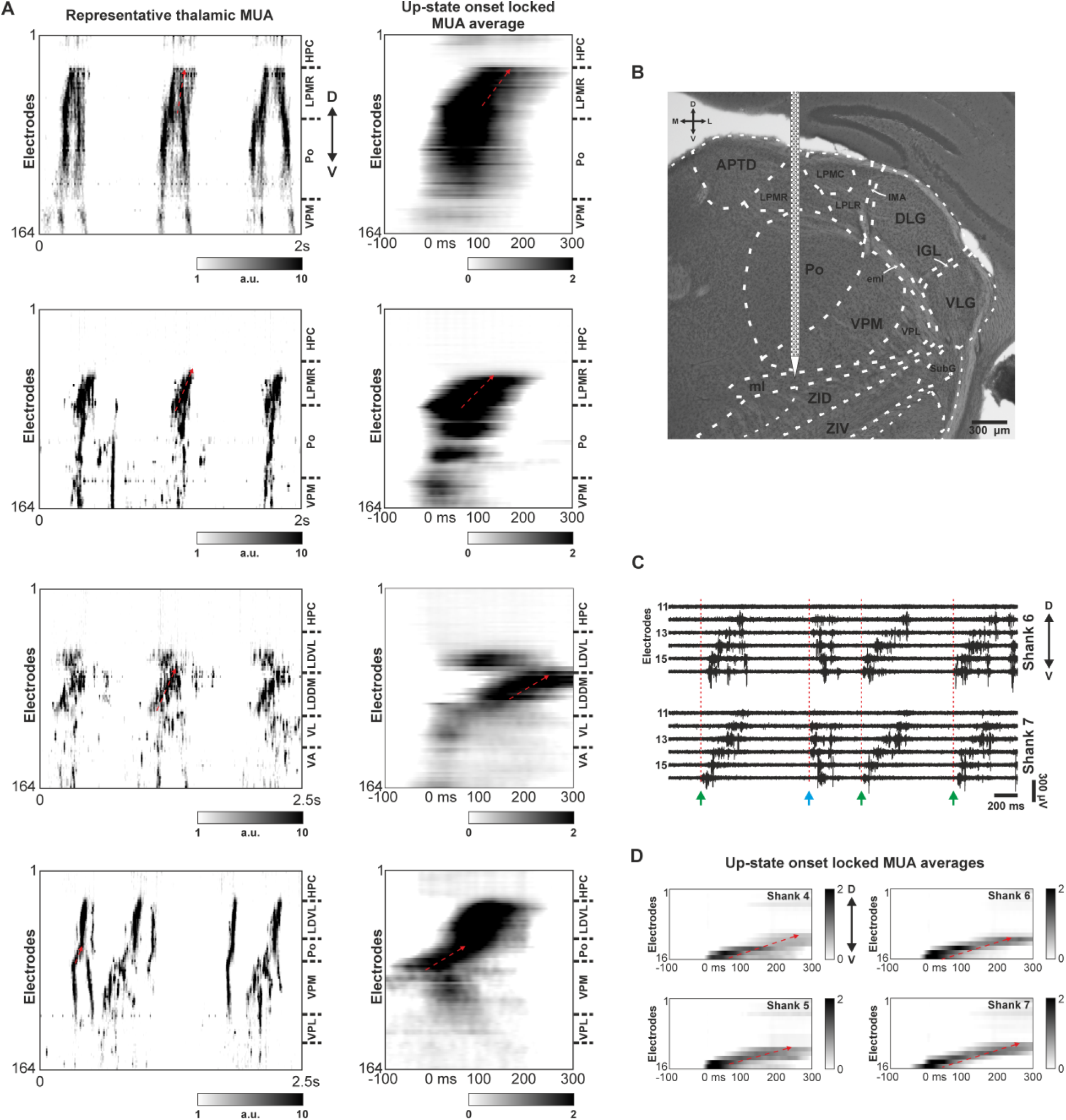
Multiunit activity propagation during up-states observed in the dorsal thalamus of ketamine/xylazine-anesthetized mice. (A) Left: Two-second-long representative thalamic MUA recordings (Neuropixels) from two mice (n = 2 insertions/mice; top two rows correspond to one animal, bottom two rows correspond to the other mouse). Right: Normalized (z-score) up-state onset locked thalamic MUA depth profile averages (n = 1555, 2261, 2450, 994 up-states from top to bottom, respectively). (B) Nissl-stained coronal brain section illustrating the approximate location of the probe shank and recording sites (electrodes) relative to major thalamic nuclei. The brain section corresponds to the recording shown in the first row of panel A. (C) Example thalamic MUA recorded in a mouse on two adjacent shanks of a multi-shank probe. Up-state onsets are indicated by vertical dashed red lines. Note ventral-to-dorsal (green arrows) and dorsal-to-ventral (blue arrow) propagation patterns. (D) Normalized (z-score) MUA depth profile averages locked to the up-state onsets (n = 2132 up-states) on shanks 4-7 showing ventral-to-dorsal propagation (dashed red arrow). The MUA average was computed from the recording partially shown in panel C. In the MUA depth profiles, the up-state starts at time point zero. D – dorsal; V – ventral; M – medial; L – lateral.

### 3.5. Thalamic activity propagation is less frequent and faster during natural sleep in rats

Although thalamocortical activity induced by ketamine/xylazine is a suitable and commonly used model of neuronal activity emerging during natural slow-wave sleep (SWS; Crunelli and Hughes, 2010; Neske, 2016), these artificially generated slow waves differ in some of their properties compared to slow waves occurring in natural sleep (Chauvette et al., 2011). To examine whether the properties of thalamic MUA propagation differ between anesthesia-induced slow waves and slow waves emerging during SWS, we chronically implanted Neuropixels probes in the thalamus of two rats, targeting thalamic nuclei where MUA propagation was most frequently observed (i.e., Po and LP). After the animals recovered from the implantation surgery, we recorded their sleep-wave cycles on a daily basis for several days, then identified sleep and wake periods in these recording sessions (Supplementary Fig. S1). Thalamic activity during wake periods and rapid eye movement (REM) sleep stages was desynchronized with MUA of active neuronal populations showing no signs of propagation (Fig. 9A). Furthermore, discrimination of thalamic nuclei based on thalamic MUA was not possible during wake and REM periods. In contrast, during non-rapid eye movement (NREM) sleep, thalamic population activity was usually synchronized, and MUA propagation could be detected sporadically (Fig. 9A). Moreover, during NREM sleep, adjacent thalamic nuclei could be more easily distinguished based on the recorded population activity (Fig. 9). Since neuronal synchronization was not as high as under ketamine/xylazine anesthesia, neuronal activity in the examined thalamic nuclei was independent from each other most of the time (e.g., between VPM and Po/LP) displaying short MUA events which were localized to a single nucleus and showed different levels of synchronization. However, for short time periods (i.e., for the duration of several subsequent up-states), MUA events between adjacent nuclei became synchronized with a small subset of these events also exhibiting propagation.

**Figure 9.**
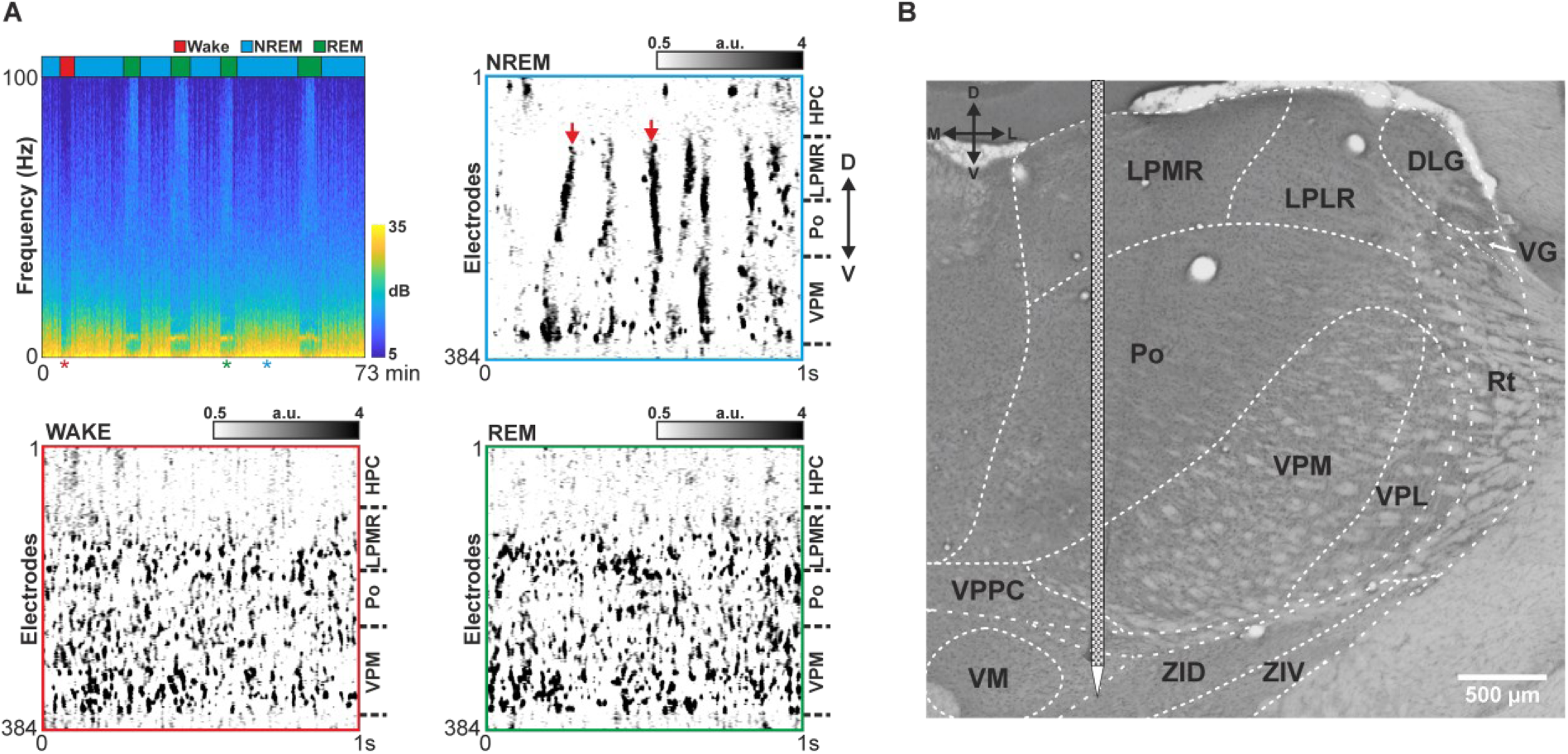
Thalamic MUA propagation is present only during non-rapid eye movement (NREM) sleep in naturally sleeping rats. (A) Spectrogram (top left) of a 73-minute-long recording session from a chronically implanted (Neuropixels probe) rat showing multiple periods of NREM sleep, REM (rapid eye movement) sleep and wake. A short thalamic MUA snippet is shown for each sleep/wake state (top right, NREM; bottom left, wake; bottom right, REM). Red arrows in the NREM sleep recording demonstrate examples of propagating population activity in the thalamus. Asterisks below the spectrogram mark the time points of the MUA snippets. The color-coded horizontal bars above the spectrogram indicate sleep and wake periods. Boundaries of major thalamic nuclei are marked by horizontal dashed lines on the right side of the MUA maps. (B) Nissl-stained coronal brain section illustrating the approximate location of the probe shank and recording sites (electrodes) relative to major thalamic nuclei in one of the chronically implanted rats.

Although we were able to detect the propagation of thalamic population activity during NREM sleep, the rate of occurrence of these events was much lower (1.89 ± 0.72 events/min) compared to ketamine/xylazine anesthesia where clear propagation was observed, on average, during every second up-state (∼45 events/min considering 1.5 Hz slow oscillation frequency). To directly compare the properties of propagating thalamic events between anesthesia and natural SWS, we computed the average thalamic MUA depth profiles locked to propagating MUA events visually detected during NREM periods, as well as MUA depth profiles locked to up-state onsets detected during light anesthesia (Fig. 10). Qualitative properties of thalamic up-states emerging during light anesthesia were very similar to those recorded during deep anesthesia, including the preferred direction of propagation (e.g., compare Fig. 4A vs. Fig. 10A). Analogous to anesthesia-induced up-states, thalamic propagating MUA events detected during NREM sleep also displayed a ventral-to-dorsal spread of activity (Fig. 10A vs. Fig. 10B). However, these MUA events were shorter, and the activity propagation was faster compared to up-states recorded under ketamine/xylazine anesthesia (Fig. 10A vs. Fig. 10B). Interestingly, thalamic MUA depth profiles were very similar between the two rats examined, suggesting that consistent and stereotyped activity patterns emerge in this part of the thalamus both during natural sleep and anesthesia (Fig. 10A1 vs. Fig. 10A2; Fig. 10B1 vs. Fig. 10B2).

**Figure 10.**
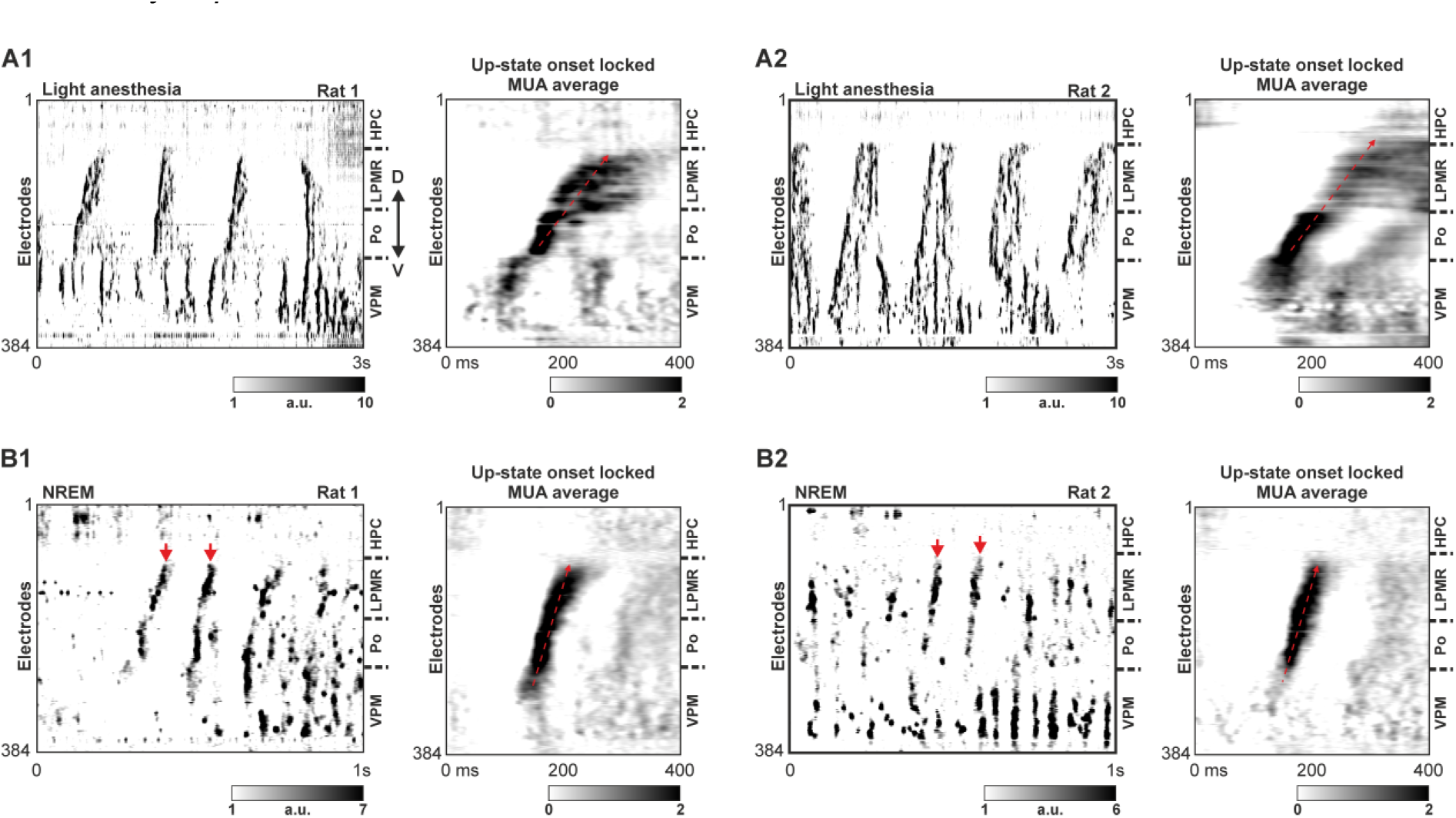
Propagating thalamic population activity during light anesthesia and natural NREM sleep. (A1-A2) Left: representative thalamic MUA recordings obtained under light anesthesia with a Neuropixels probe at the end of the implantation surgery in two chronically implanted rats. Right: Normalized (z-score) up-state onset locked thalamic MUA depth profile averages (n = 90 and 589 up-states for the two rats, respectively). (B1-B2) Left: representative MUA recordings with propagating activity obtained during NREM sleep in the two chronically implanted animals. Right: Normalized (z-score) thalamic MUA depth profile averages locked to the onset of detected propagating MUA events (n = 407 and 141 up-states for the two rats, respectively). Red arrows in the NREM recording mark examples of propagation population activity in the thalamus. Boundaries of major thalamic nuclei are indicated by horizontal dashed lines on the right side of the MUA maps. Oblique dashed red arrows indicate the direction of propagation. Note the difference in propagation speed between anesthesia-induced MUA propagation and MUA propagation during NREM sleep. D – dorsal, V – ventral.

## 4. Discussion

The present work demonstrates, for the first time, that propagating patterns of spontaneous population activity occur in the rodent thalamus in vivo, both under anesthesia and during slow-wave sleep. Additionally, our findings revealed that neuronal activity is most likely to propagate within dorsal, particularly higher order thalamic nuclei, where up-states generally travel along the dorsoventral axis with a preference for the ventral-to-dorsal direction. While, in general, the thalamus exhibited a rich variety of spatiotemporal activity patterns both during natural sleep and anesthesia, thalamic neuronal activity in anesthetized animals was less complex and more stereotyped. This difference in complexity is indicated, for instance, by our observation that the majority of up-states occurring under anesthesia display ventral-to-dorsal propagation with minimal variations in their propagation trajectories (Fig. 5). During natural sleep, although still present, ventral-to-dorsal propagation was less frequent, and the spreading of MUA was faster compared to thalamic activity propagation under anesthesia (Fig. 10). Furthermore, ketamine/xylazine anesthesia-induced cortical population activity, obtained with multi-shank silicon probes, also exhibited stereotypical propagation patterns. A notable proportion of cortical up-states arrived first at the anterior shanks, then traveled in the posterior direction (Figs. 6 and 7). The proportion of these anterior-to-posterior propagating cortical up-states was similar to the proportion of predominant ventral-to-dorsal propagating thalamic up-states (∼50% of up-states), whereas proportions of other propagation patterns were notably lower (3-10%; Fig. 7A and B). These observations are in line with the recent theory that the strongly interconnected neocortex and thalamus function as a single slow wave generating unit (Crunelli et al., 2015). However, ventral-to-dorsal propagation of thalamic activity was more pronounced in thalamic MUA depth profiles locked to cortical up-states than in those aligned to thalamic up-states (Fig. 6 vs. Supplementary Fig. S7). This suggests that the neocortex may play an influential role in shaping population activity within the thalamus.

Since its discovery, numerous properties of slow wave propagation have been described in both humans and animal models. In humans, for instance, based on high-density EEG recordings, multiple studies have confirmed that individual spontaneous slow waves emerging during NREM sleep can basically start at any site of the cortex and then typically travel along the anterior-posterior axis with unique trajectories (Massimini et al., 2004; Murphy et al., 2009; Ujma et al., 2022). While these global slow waves can start anywhere within the cortex, they preferentially originate in the insula or the cingulate gyrus. From there, they propagate in the anterior-to-posterior direction, terminating in parietal or occipital cortical areas (Massimini et al., 2004; Murphy et al., 2009). Spontaneous slow waves in humans travel at an average speed of 2.2 m/s with propagation delays ranging from 40 to 360 ms (Massimini et al., 2004; Murphy et al., 2009). Using ECoG recordings, Hangya and colleagues investigated the spatiotemporal dynamics of slow wave propagation on a finer spatial scale (∼1 cm; Hangya et al., 2011). They found that, compared to the propagation of global slow waves observed in EEG recordings, localized slow waves exhibit more complex spatiotemporal dynamics, displaying a diverse set of propagation patterns, including convergence, divergence, or circular patterns (Hangya et al., 2011).

In rodents, propagation of spontaneous cortical slow waves has been revealed using a diverse array of experimental techniques, including voltage-sensitive dye imaging, two-photon and wide-field calcium imaging, as well as multielectrode and micro-ECoG recordings (Dasilva et al., 2021; Ferezou et al., 2007; Fucke et al., 2011; Greenberg et al., 2018; Greenberg and Dickson, 2013; Luczak et al., 2007; Matsui et al., 2016; Reyes-Puerta et al., 2016; Ruiz-Mejias et al., 2011; Sheroziya and Timofeev, 2014; Stroh et al., 2013; Takagaki et al., 2008; Vyazovskiy et al., 2009; Xu et al., 2007). These in vivo studies have investigated the properties of slow waves in both naturally sleeping and anesthetized animals. In the anesthetized models, a wide variety of anesthetics have been used to induce slow waves, including urethane, ketamine/xylazine, isoflurane, and chloral hydrate. In rats, similarly to humans, spontaneous global slow waves occurring in the neocortex had a tendency to propagate along the anterior-posterior axis both under anesthesia (Greenberg and Dickson, 2013; Takagaki et al., 2008) and during NREM sleep (Vyazovskiy et al., 2009). However, slow waves examined in localized cortical areas could exhibit distinct propagation patterns (e.g., propagation along the medial-lateral axis in the visual cortex; Wanger et al., 2013) or did not show a preferred propagation direction (Luczak et al., 2007; Reyes-Puerta et al., 2016; Xu et al., 2007). In mice, similarly to rats and humans, most studies found that cortical slow waves preferentially travel along the anterior-to-posterior axis (Brier et al., 2019; Dasilva et al., 2021; Greenberg et al., 2018; Liang et al., 2021; Matsui et al., 2016; Niethard et al., 2018; Ruiz-Mejias et al., 2011; Sheroziya and Timofeev, 2014; Stroh et al., 2013; Vanni et al., 2017), predominantly propagating in the anterior-to-posterior direction (Brier et al., 2019; Greenberg et al., 2018; Matsui et al., 2016; Niethard et al., 2018; Sheroziya and Timofeev, 2014; Stroh et al., 2013). In contrast, one study observed slow wave propagation in the visual cortex of mice along the medial-lateral axis (Ruiz-Mejias et al., 2011).

In rodents, the propagation speed of in vivo cortical slow waves was notably slower than in humans, falling within the range of 10 to 100 mm/s (Dasilva et al., 2021; Ferezou et al., 2006; Fucke et al., 2011; Greenberg et al., 2018; Greenberg and Dickson, 2013; Han et al., 2008; Reyes-Puerta et al., 2016; Ruiz-Mejias et al., 2011; Sheroziya and Timofeev, 2014; Stroh et al., 2013; Takagaki et al., 2008; Wanger et al., 2013), whereas propagation of cortical population activity was somewhat slower in vitro (5-10 mm/s; Reig et al., 2010; Sanchez-Vives and McCormick, 2000). Our findings related to cortical slow wave propagation are in good agreement with the results above. Even within a localized cortical area (∼1.5 mm), we were able to observe a tendency for up-states to travel in the anterior-to-posterior direction with a propagation speed (∼38 mm/s) within the range reported in earlier studies. Interestingly, in the study by Reyes-Puerta and colleagues, who examined slow waves in urethane-anesthetized rats using voltage-sensitive dye imaging and silicon probe recordings in cortical areas of similar extent (∼1.5 mm), the observed spatiotemporal pattern of cortical up-state propagation closely resembled our data (compare Fig. 7C and D vs. Fig. 1E and F in Reyes-Puerta et al., 2016). They found that the average propagation speed of spontaneous cortical slow waves is about 35 mm/s, very close to the average speed measured in our cortical recordings.

There are significantly fewer studies investigating the spatiotemporal dynamics of slow waves outside the cortex. In rats anesthetized with chloral hydrate, Ushimaru and colleagues found that neurons in ventral anterior and ventromedial (VA/VM) thalamic nuclei, as well as in the reticular thalamic (Rt) nucleus, fire notably earlier (approximately at the start of up-states) compared to neurons located in the VL nucleus and layer 5 of the frontal cortex (Ushimaru et al., 2012). Furthermore, in ketamine/xylazine-anesthetized rats, Slezia and colleagues found that thalamocortical neurons located in different thalamic nuclei (VPM, VPL and Po) fire at distinct phases of up-states (Slezia et al., 2011). For instance, population activity in the Po preceded firing in VPM/VPL nuclei with a time lag of about 100 ms. Our findings are consistent with the delay observed in these nuclei. In our thalamic recordings, population activity in the ventral parts of the Po started on average several tens of milliseconds earlier than MUA in the VPM (see Fig. 6). The leading role of Po in up-state generation is further supported by the study of Sheroziya and Timofeev (Sheroziya and Timofeev, 2014), where they found that in ketamine/xylazine anesthetized mice, the Po was among the first structures to fire during up-states, along with frontal and motor cortices, and the parafascicular (PF) thalamic nucleus. Thalamocortical structures with the latest activity were the posterior cortex and the anteroventral (AV) thalamic nucleus. In our simultaneous cortical and thalamic recordings, in both rats and mice, we observed that MUA in ventral parts of the Po tended to start before population activity during cortical up-states and also preceded MUA in adjacent thalamic nuclei (see Figs. 6 and 8). Interestingly, a human study using EEG and depth electrode recordings found only short time lags between spontaneous cortical and thalamic slow waves in pharmacoresistant epileptic patients (Ujma et al., 2022). Thalamic slow waves started nearly simultaneously in all examined thalamic nuclei (i.e., the anterior thalamus, mediodorsal and ventral anterior nuclei), shortly after cortical slow waves. While methods based on field potentials might be less precise in determining propagation patterns of smaller neuronal populations comparted to MUA-based methods, the study suggests that, similar to rodents, the propagation of slow waves does not occur in every region of the thalamus.

What could potentially explain the observed preference for ventral-to-dorsal propagation in the thalamus? Most of the dorsal thalamic nuclei investigated in this study (e.g., Po, LD or LP nuclei) are classified as higher order nuclei that receive abundant input connections from pyramidal cells located in layer 5 of the neocortex (Sherman, 2007). For example, in the Po, thalamocortical neurons are strongly innervated by giant excitatory terminals originating in layer 5 of the somatosensory cortex (Bourassa et al., 1995; Hoogland et al., 1987; Veinante et al., 2000). A multitude of in vivo and in vitro animal studies revealed the pivotal role of neurons located in cortical layer 5 in the initiation and propagation dynamics of cortical up-states (Beltramo et al., 2013; Chauvette et al., 2010; Fiath et al., 2016b; Sakata and Harris, 2009; Sanchez-Vives and McCormick, 2000; Stroh et al., 2013; Wester and Contreras, 2012). Furthermore, it is well established that sensory cortices, such as the primary somatosensory or visual cortex, exhibit topographical organization. Since the Po primarily receives its inputs from the somatosensory cortex, whereas the more dorsally located LP nucleus is interconnected with visual cortical areas, and considering the anterior positioning of the somatosensory cortex relative to the visual cortex, it is reasonable to predict that, during up-states traveling from anterior to posterior cortical areas, MUA induced by the firing of cortical layer 5 pyramidal cells in the Po would precede neuronal activity in the LP, as was also observed in our data. Thus, we speculate that MUA propagation occurring in the thalamus is primarily a consequence of the horizontal propagation of cortical slow waves. In this scenario, thalamocortical neurons are sequentially activated via the intense firing of layer 5 pyramidal cells. Consequently, the anterior-to-posterior cortical slow wave propagation would translate to a ventral-to-dorsal propagation in the thalamus.

The above theory is supported by our observation that up-states displaying cortical anterior-to-posterior propagation were detected in comparable proportions to thalamic up-states with ventral-to-dorsal propagation (see Fig. 7). In addition, cortical up-states traveling in the opposite direction (posterior-to-anterior) were frequently observed with dorsal-to-ventral thalamic propagation (Fig. 7), providing further support for our hypothesis. Furthermore, if cortical up-state propagation is relayed to higher order nuclei via layer 5 pyramidal cells, this would partially explain the gradual propagation observed between Po and LP nuclei (e.g., Fig. 10), since somatosensory and visual cortical areas are practically adjacent to each other. For instance, an up-state traversing the somatotopic map of the somatosensory cortex towards the adjacent visual cortex through horizontal cortico-cortical connections would reach the visual cortex with minimal delay and continue its course across the retinotopic map. If layer 5 cortical inputs are topographically organized within these nuclei, this cortical propagation would appear in the thalamus as a relatively smooth MUA propagation both within and between Po and LP, aligning with our observations in thalamic recordings. Finally, the observed ∼100 ms delay between the up-state initiation and the termination of MUA propagation in the dorsal thalamus closely mirrors the ∼100 ms time lag of slow waves found between the frontal and posterior areas of the neocortex (Stroh et al., 2013). The similarity in time lags between the cortex and the thalamus implies that the propagation speed of cortical slow waves must be higher than that of thalamic slow waves, given that up-states have to travel greater distances across the cortical mantle than thalamic up-states between thalamic nuclei. In agreement with this assumption, in our thalamocortical recordings, we observed a more than three times faster propagation of cortical up-states compared to propagation of thalamic up-states.

In several of our thalamic recordings, we noticed that population activity in adjacent higher order thalamic nuclei (e.g., Po and LD) may decouple, that is, the gradual MUA propagation between these areas was absent, and MUA propagated separately within each nucleus, or no MUA propagation was observed at all. This decoupling phenomenon was more frequent under lighter anesthesia or during slow-wave sleep. As observed in prior studies, these states are characterized by increased variability and complexity in cortical propagation patterns (Dasilva et al., 2021; Pazienti et al., 2022). These studies demonstrated that lighter anesthesia leads to an increased spatiotemporal complexity of cortical activity, manifesting in a higher number of distinct propagation patterns/modes (exhibiting more diverse spatiotemporal patterns) and an increased propagation speed compared to deeper levels of anesthesia (Dasilva et al., 2021; Pazienti et al., 2022). Similar results were found for the transition from anesthetized states to wakefulness (Liang et al., 2023). In our study, we also observed an increase in the number of distinct thalamic MUA patterns and in the speed of MUA propagation between light anesthesia and slow-wave sleep (Fig. 10).

While our observations provide strong evidence for the occurrence of spontaneous activity propagation in the rodent thalamus, it is important to discuss the limitations of our study. First, despite our efforts to comprehensively explore the activity of the thalamus, not all thalamic nuclei were investigated here. For example, we did not record the activity of midline and intralaminar thalamic nuclei, due to their complicated access with silicon probes, irregular geometries, and smaller volumes relative to the nuclei we examined. These factors would complicate the targeting and identification of these nuclei, and thus the reliable detection of MUA propagation. However, we can not rule out the possibility that propagation of thalamic population activity might also occur in these nuclei. Second, we did not perform spike sorting and single-unit activity analysis in this study. Nevertheless, we would not expect any qualitative change in our results by analyzing single-unit activity instead of MUA, since propagation of thalamic up-states could be reliably identified based on the population activity alone. However, an in-depth analysis of thalamic activity on a single unit level could yield deeper insights into the properties of thalamic propagation patterns in future investigations. Third, cortical population activity was obtained only from a local cortical area (∼1.5 mm along the anterior-posterior axis) which limits our field of view to observe cortical up-state propagation. Extending our observation by involving a larger cortical area (e.g., by implanting multiple silicon probes into the cortex), would enable a more comprehensive examination of the interplay between cortical and thalamic population activity. Finally, in some of our thalamic recordings from anesthetized rats we noticed that neuronal activity patterns might change over time (suddenly or over longer time periods), transitioning from exhibiting MUA propagation (indicating a state with higher neuronal synchronization) to displaying no propagation in the same thalamic nuclei (reflecting a less synchronized state), and vice versa. This implies that the actual brain state (e.g., the depth of anesthesia) could also have a strong impact on ongoing thalamic activity patterns, and that MUA propagation might occur even in more thalamic nuclei than actually observed, since a consistent depth of ketamine/xylazine anesthesia is challenging to maintain.

## Acknowledgements

Project no. PD134196 (RF) has been implemented with the support provided by the Ministry of Culture and Innovation of Hungary from the National Research, Development and Innovation Fund, financed under the PD_20 funding scheme. RF was supported by the Bolyai János Scholarship of the Hungarian Academy of Sciences. The research leading to these results has received funding from the Hungarian Brain Research Program Grant (2017-1.2.1-NKP-2017-00002, NAP2022-I-2/2022) and the Pharmaceutical Research and Development Laboratory project (PharmaLab, RRF-2.3.1-21-2022-00015). Archiving of research data was supported by the ARP (Data Repository Platform) project of the Eötvös Loránd Research Network.

## CRediT authorship contribution statement

Csaba Horváth: Methodology, Formal analysis, Investigation, Data Curation, Visualization, Writing - Original Draft, Writing – Review & Editing. István Ulbert: Conceptualization, Writing - Original Draft, Writing – Review & Editing, Supervision, Funding acquisition. Richárd Fiáth: Conceptualization, Methodology, Software, Formal analysis, Investigation, Data Curation, Validation, Visualization, Writing - Original Draft, Writing – Review & Editing, Project administration, Supervision, Funding acquisition.

