## Supplementary Material for "Propagating population activity patterns during spontaneous slow waves in the thalamus of rodents"

| Abbreviation | Brain region |
|---|---|
| AM | anteromedial nucleus |
| APT | anterior pretectal nucleus |
| APTD | dorsal part of the anterior pretectal nucleus |
| AV | anteroventral nucleus |
| AVDM | dorsomedial part of the anterovent nucleus |
| bsc | brachium of the superior colliculus |
| CL | centrolateral nucleus |
| DLG | dorsal lateral geniculate nucleus |
| eml | external medullary lamina |
| EpP | epipeduncular nucleus |
| HPC | hippocampus |
| ic | internal capsule |
| IGL | intergeniculate leaf |
| IMA | intramedullary thalamic area |
| InG | intermediate gray layer of the superior colliculus |
| InWh | intermediate white layer of the superior colliculus |
| LD | lateral dorsal nucleus |
| LDDM | dorsomedial part of the laterodorsal nucleus |
| LDVL | ventrolateral part of the laterodorsal nucleus |
| LPLR | laterorostral part of the lateral posterior nucleus |
| LPMC | mediocaudal part of the lateral posterior nucleus |
| LPMR | mediorostral part of the lateral posterior nucleus |
| MGD | dorsal part of the medial geniculate nucleus |
| MGM | medial part of the medial geniculate nucleus |
| MGV | ventral part of the medial geniculate nucleus |
| ml | medial lemniscus |
| MZMG | marginal zone of the medial geniculate |
| OT | nucleus of the optic tract |
| PaR | pararubral nucleus |
| PLi | posterior limitans nucleus |
| PoT | triangular part of the posterior nuclear group |
| PP | peripeduncular nucleus |
| Rt | reticular thalamic nucleus |
| SG | suprageniculate thalamic nucleus |
| SNL | lateral part of the substantia nigra |
| str | superior thalamic radiation |
| SubG | subgeniculate nucleus |
| VA | ventral anterior nucleus |
| VG | ventral geniculate nucleus |
| VL | ventrolateral nucleus |
| VLG | ventral lateral geniculate nucleus |
| VLi | ventral linear nucleus of the thalamus |
| VM | ventromedial nucleus |

| | |
|---|---|
| VPL | ventral posterolateral nucleus |
| VPM | ventral posteromedial nucleus |
| VPPC | parvicellular part of the ventral posterior nucleus |
| ZIC | caudal part of the zona incerta |
| ZID | doral part of the zona incerta |
| ZIV | ventral part of the zone incerta |

**Supplementary Table S1.** List of abbreviations of brain regions used in the text and figures.

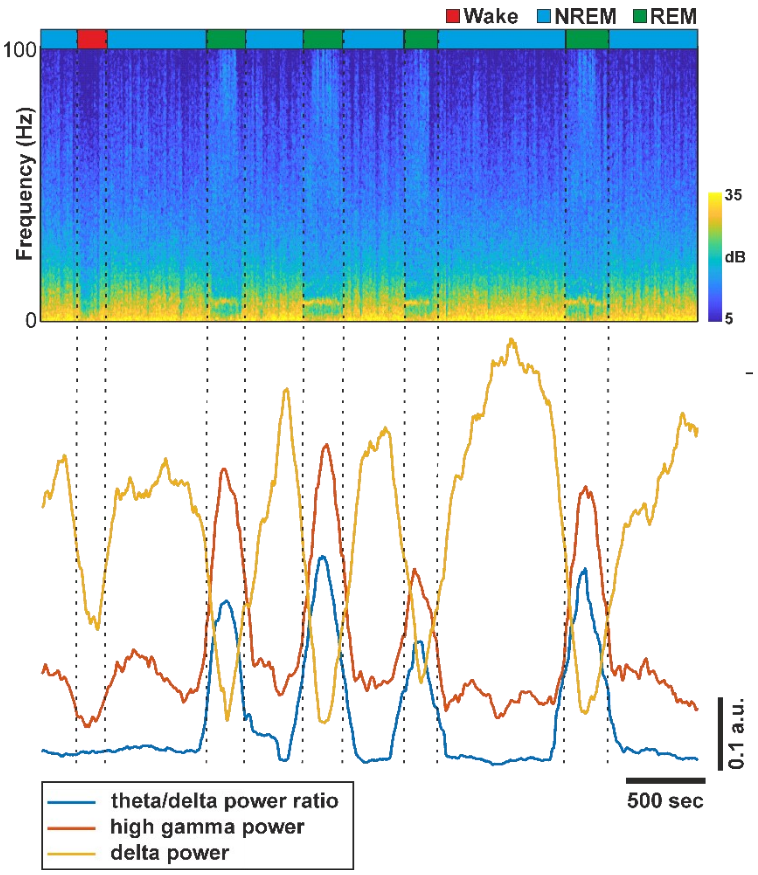


**Supplementary Figure S1.** Example of sleep stage scoring in chronically implanted, naturally sleeping rats. The time course of several spectral features (e.g., power in the delta frequency band) were calculated from the spectrogram (top) and used to determine the actual sleep stage (see the Methods section for details). Vertical dashed black lines indicate the transition between different sleep stages. NREM – non-rapid eye movement sleep, REM – rapid eye movement sleep.

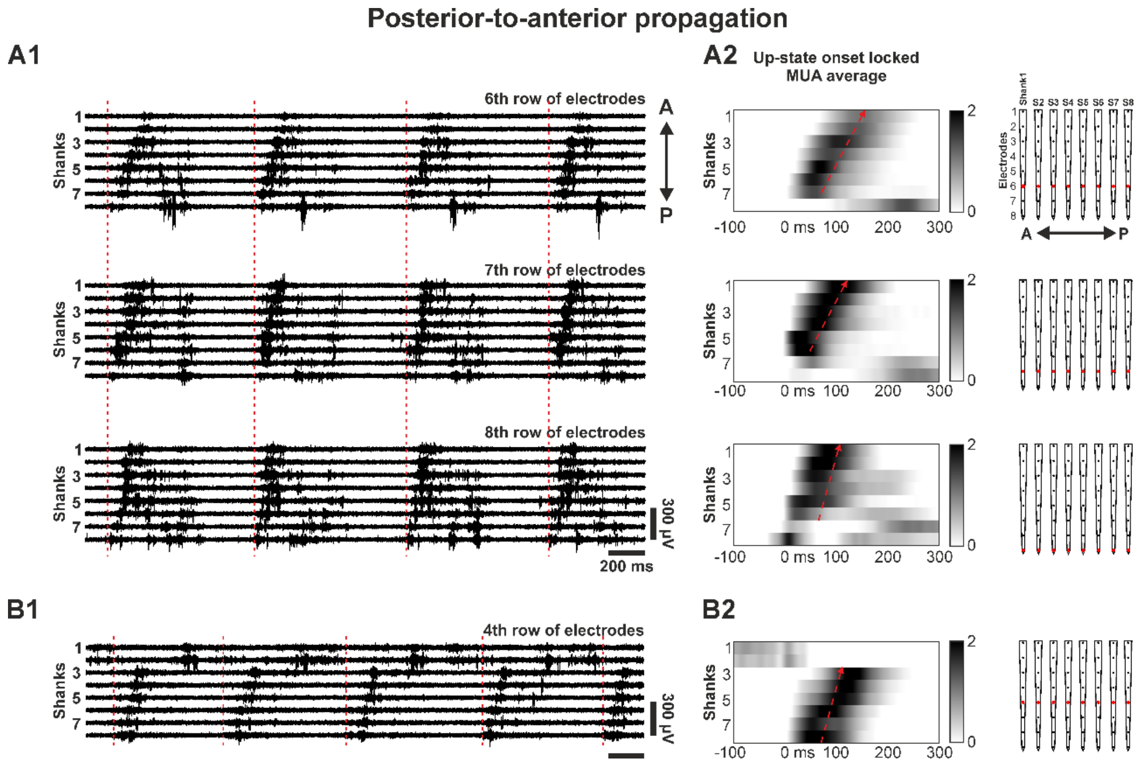


**Supplementary Figure S2.** Posterior-to-anterior propagation of population activity observed with multi-shank silicon probes in the thalamus of anesthetized rats. (A1) Representative thalamic multiunit activity (MUA) recording obtained by three adjacent rows of electrodes on eight shanks of the recording probe from an anesthetized rat. Up-state onsets are indicated by dashed red vertical lines. (A2) Normalized (z-score) MUA depth profile averages locked to the up-state onsets (n = 2007 up-states) on the three rows of electrodes showing posterior-to-anterior propagation in the thalamus (oblique dashed red arrow). The up-state starts at time point zero. The row of electrodes used is indicated on the right in red. (B1-B2) Example recording from another rat obtained with a single row of electrodes (n = 2051 up-states). A – anterior; P – posterior.

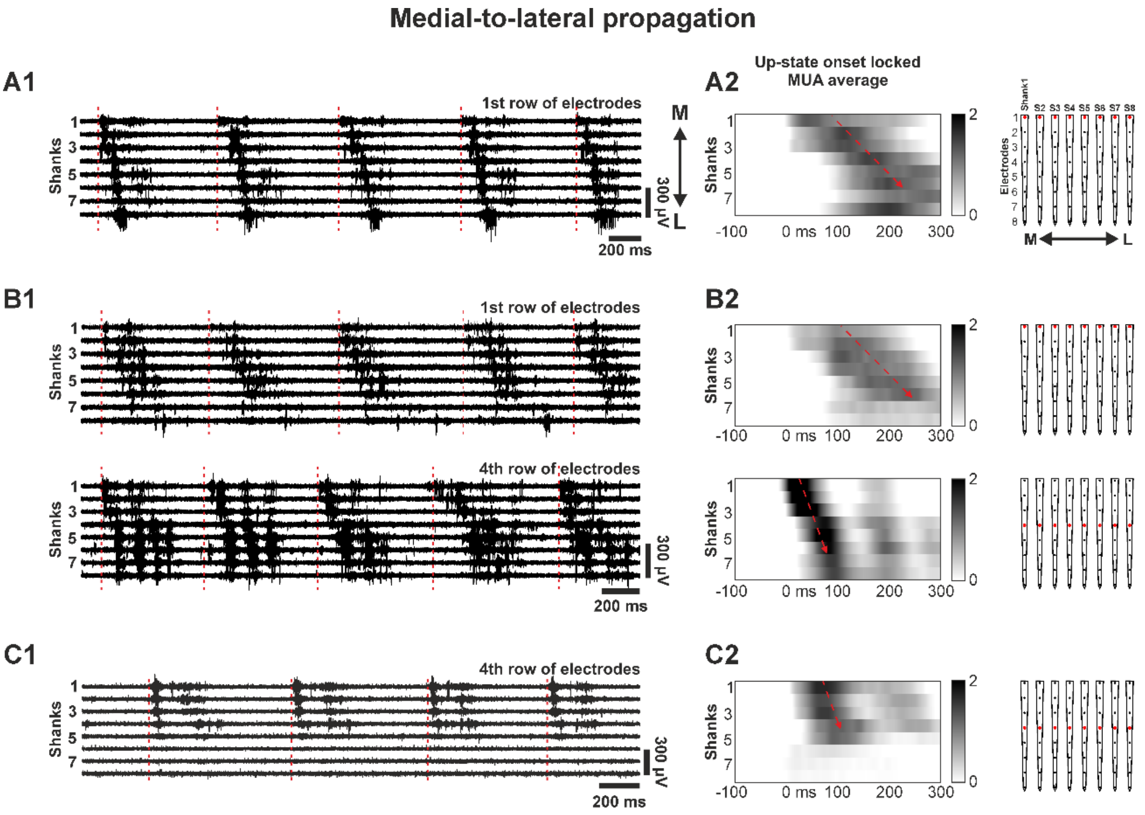


**Supplementary Figure S3.** Medial-to-lateral propagation of population activity observed with multi-shank silicon probes in the thalamus of anesthetized rats. (A1) Representative thalamic MUA recording obtained by a single row of electrodes on eight shanks of the recording probe from an anesthetized rat. Up-state onsets are indicated by dashed red vertical lines. (A2) Normalized (z-score) MUA depth profile average locked to the up-state onsets (n = 2608 up-states) on the row of electrodes showing medial-to-lateral propagation in the thalamus (oblique dashed red arrow). The up-state starts at time point zero. The row of electrodes used is indicated on the right in red. (B1-B2 and C1-C2) Example recordings from two other rats (n = 1983 and 1867 up-states, respectively). M – medial; L – lateral.

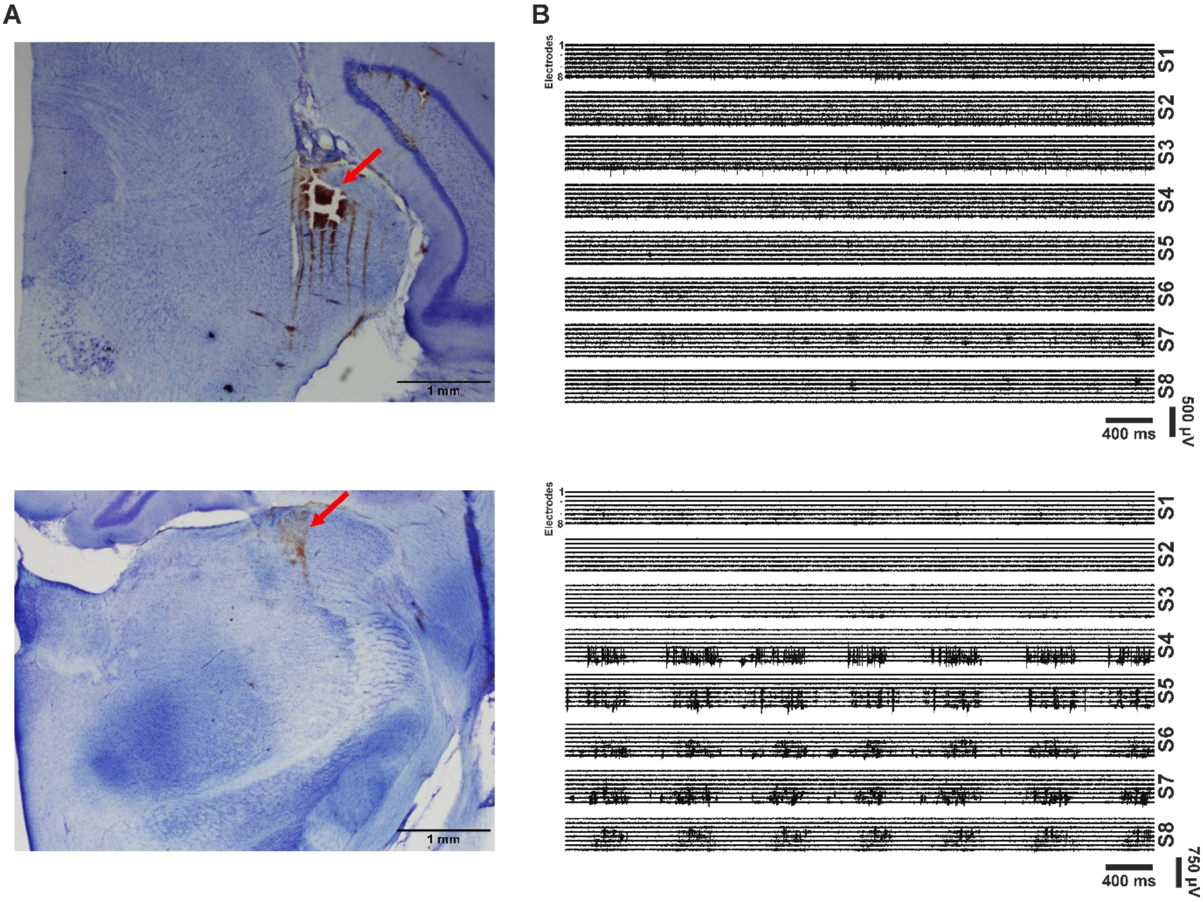


**Supplementary Figure S4.** (A) Nissl-stained coronal rat brain slices with signs of bleeding and tissue damage in the thalamus (red arrows). (B) Sample MUA traces recorded with multi-shank probes at thalamic sites shown in panel A. Note the significantly reduced MUA on all shanks (S1-S8) in the top example and on the first three shanks (S1-S3) in the bottom example.

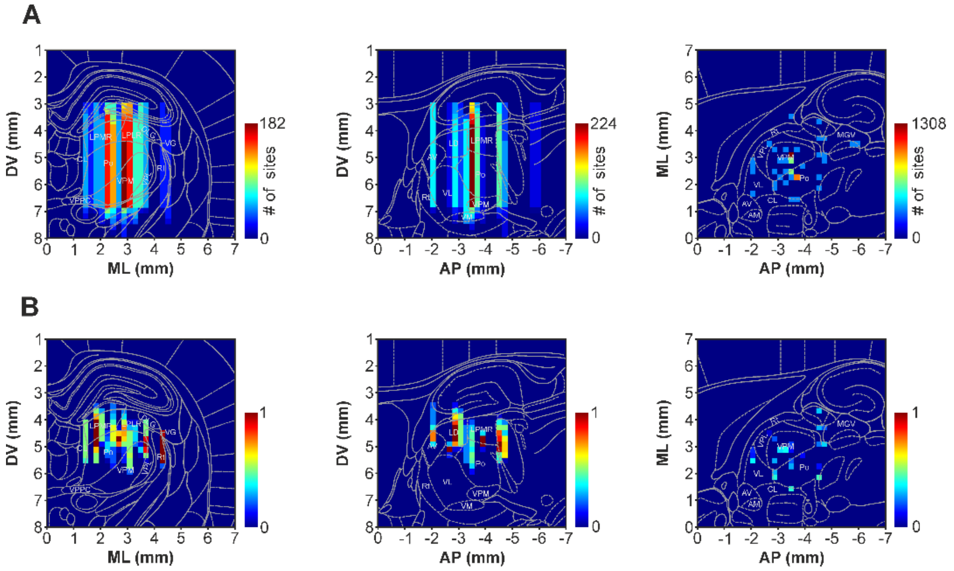


**Supplementary Figure S5.** Colormaps showing the distribution of all recording site locations used to collect thalamic activity with single-shank Neuropixels high-density silicon probes in anesthetized rats (A) along the three anatomical axes (left: anterior-posterior (AP) axis; middle: medial-lateral (ML) axis; right: dorsal-ventral (DV) axis), and the probability of detecting propagating activity along the dorsoventral axis at these sites (B). Each pixel within these colormaps represents a small quadratic region in the brain with an area of 0.2 mm x 0.2 mm. The pixel values represent cumulative values along one of the three anatomical axes. Schematic brain sections overlaid on the colormaps indicate the location of relevant thalamic nuclei at coordinates AP -3.6 mm (left), ML 1.9 mm (middle) and DV 5.6 mm (right), relative to the bregma.

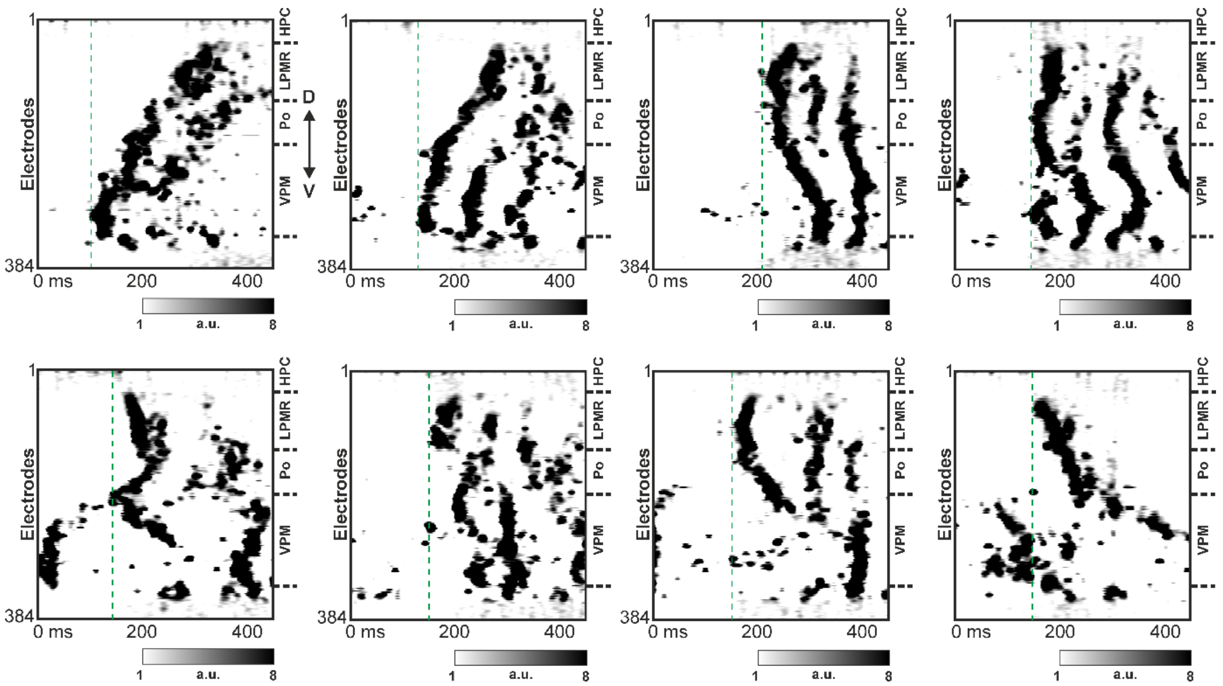


**Supplementary Figure S6.** Examples showing the variety of thalamic propagation patterns in a single recording from an anesthetized rat. Boundaries of major thalamic nuclei are marked by horizontal dashed lines on the right side of the multiunit activity maps. Vertical dashed green lines mark the approximate onset of up-states.

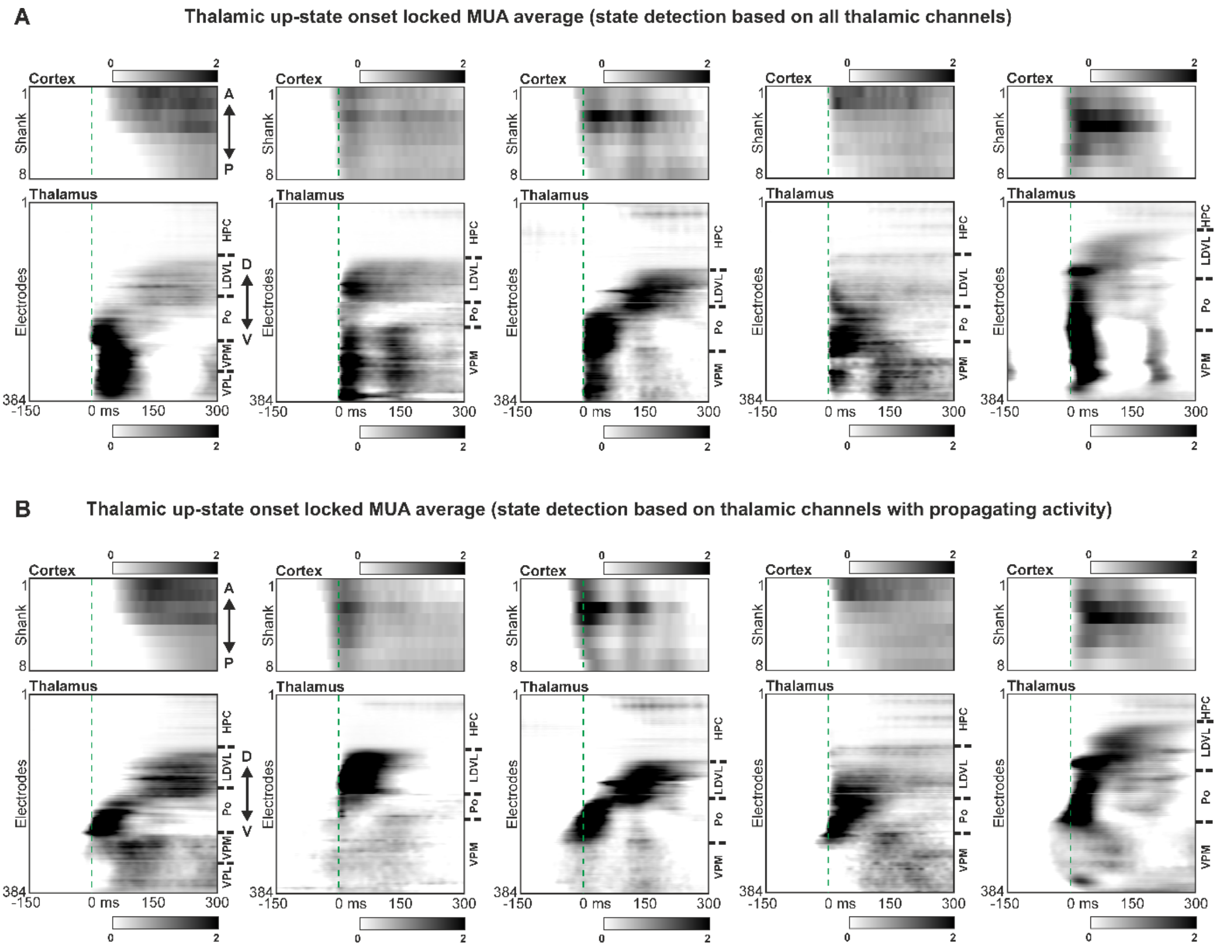


**Supplementary Figure S7.** (A) Normalized (z-score) thalamic up-state onset locked (using all thalamic channels for up-state onset detection) cortical (top) and thalamic (bottom) MUA depth profile averages of simultaneous cortical and thalamic recordings from five different animals (n = 4721, 4459, 2215, 2469 and 3577 up-states from left to right, respectively). (B) Here, only that part of the thalamus was selected for up-state detection that showed activity propagation (n = 3000, 3419, 1792, 2069 and 4180 up-states from left to right, respectively). Boundaries of major thalamic nuclei are marked by horizontal dashed lines on the right side of the MUA maps. Vertical dashed green lines mark the thalamic up-state onsets.
